# GRHL and PGR control WNT4 expression in the mammary gland via 3D looping of conserved and species-specific enhancers

**DOI:** 10.1101/2025.04.11.648333

**Authors:** Yorick BC van de Grift, Marleen T Aarts, Katrin E Wiese, Nika Heijmans, Ingeborg B Hooijkaas, Colin EJ Pritchard, Linda Henneman, Paul JA Krimpenfort, Antonius L van Boxtel, Renée van Amerongen

## Abstract

WNT4 is critical for epithelial side branching in the mammary gland. In both mice and humans, its expression is tightly regulated and restricted to hormone-responsive, mature luminal cells. Although generally assumed to act downstream of progesterone, the molecular mechanisms that control lineage-specific induction of *WNT4* gene expression remain unknown. Here we functionally dissect the cis-acting enhancer network and spatiotemporal transcriptional mechanisms that regulate WNT4 expression in the mouse mammary gland and the human breast. We identify Grainyhead-like (GRHL) proteins as luminal-specific pioneer factors that bind to a conserved distal chromatin hub. The progesterone receptor (PGR) binds to different cis-acting elements that contact the hub via 3D chromatin looping to induce *WNT4* expression. More generally, this model explains how the interplay between endocrine signaling and dynamic molecular interactions at the physical chromatin level can be translated into robust gene expression patterns on the tissue scale.

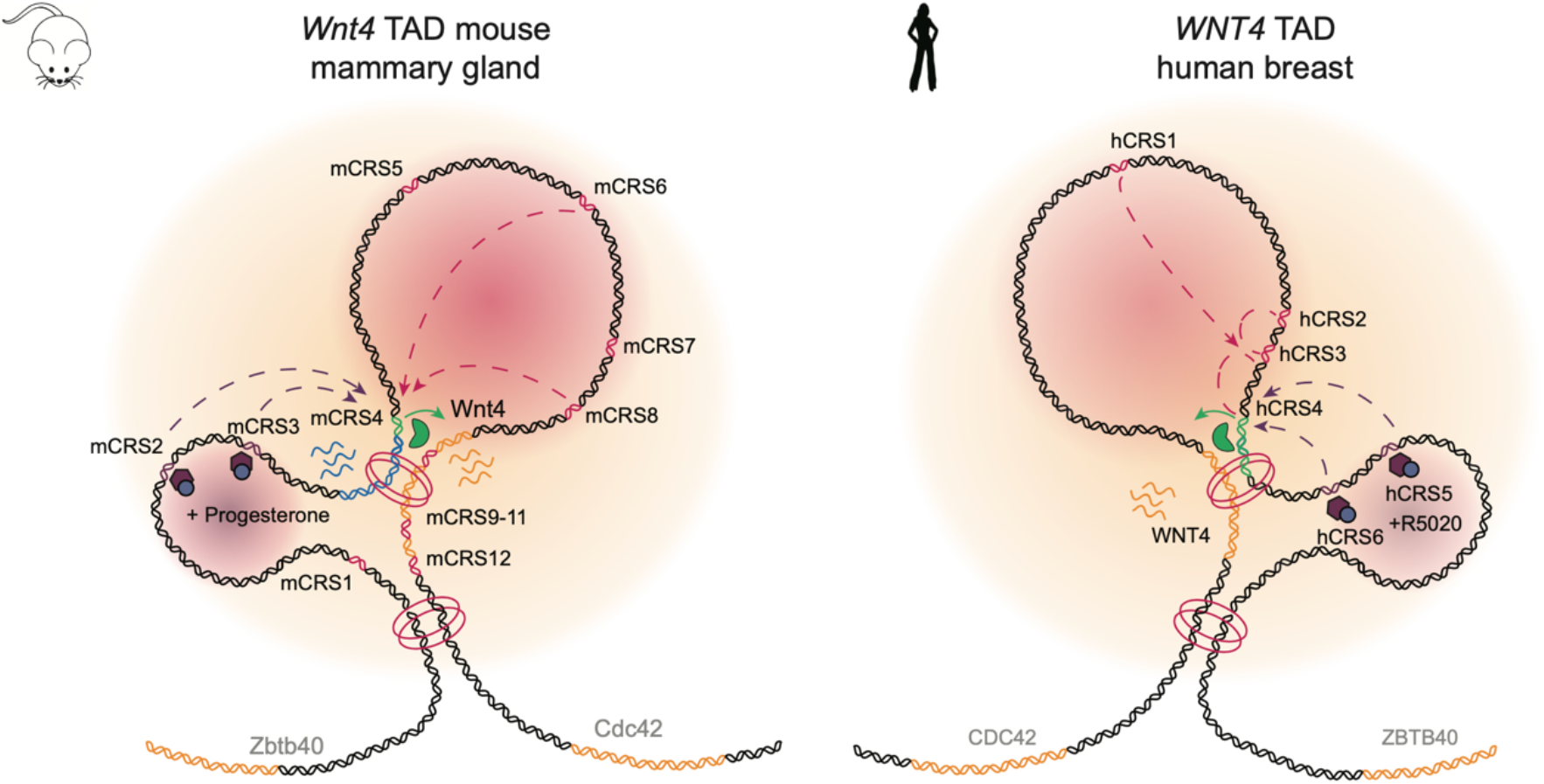

## Introduction

Multicellular animals require robust cell-cell communication mechanisms to control tissue morphogenesis and maintenance. At the molecular level this is achieved by the orchestrated activity of a surprisingly small number of evolutionarily conserved developmental signal transduction pathways. One of these is the WNT pathway, which controls fundamental biological processes throughout embryonic development and postnatal life in all metazoans, ranging from vertebrate gastrulation^1–4^ to wing color patterning in butterflies^5^. Thousands of studies have demonstrated its importance in controlling cell proliferation and cell fate decisions at different anatomical sites and life stages^6–9^. To be used in such vastly different settings, WNT signaling must simultaneously be highly flexible and highly specific, allowing different outcomes depending on the time and place of its activity. Indeed, considerable molecular complexity has evolved to elicit context-dependent signaling responses.

WNT signaling is initiated by the binding of Wnt family member (WNT) proteins to frizzled class receptor (FZD) proteins. Due to posttranslational lipid modification, WNT proteins are inherently hydrophobic^10^, which limits their diffusion in the aqueous extracellular environment. Although long-range WNT gradients have been visualized^11^ and dedicated transport mechanisms aiding their distribution exist^12–14^, WNT proteins typically act as short-range, paracrine signaling factors^15^. This means that the specificity and spatiotemporal activity of WNT signal transduction must be largely controlled at the level of *Wnt* gene expression.

The mammalian genome harbors 19 different *Wnt* genes, which show complex and dynamic expression patterns in the developing embryo^16,17^. In adult tissues, well-defined *Wnt* gene expression landscapes are maintained, providing localized sources of WNT proteins that act as self-renewal factors for somatic stem cells^18–20^. Past work has largely focused on unraveling the biochemical mechanism of WNT signal transduction, including the specificity and complexity of WNT/FZD pairing^21–24^. Attention is also increasingly being directed towards understanding the context-dependent interpretation of the WNT signal by focusing on the target gene programs that are induced in WNT-receiving cells^25,26^. Yet the most upstream initiation of the signal transduction response, namely the tissue-specific activation of *Wnt* gene expression itself, has remained virtually unexplored^27–30^. As a result, we still lack a basic understanding of the identity and function of the regulatory DNA elements and their cognate transcription factors that control this process in the context of the 3D genome.

Integrative multi-omics analyses have opened tremendous new opportunities for scientific investigation of gene regulatory mechanisms. Genome-wide approaches in particular have been used to map the global chromatin landscape and, in doing so, have revealed important organizing principles of mammalian DNA with respect to the folding, accessibility and function of its various non-coding regulatory sequences^31,32^. Despite these insights, understanding the tissue-specific regulation of any given gene remains challenging, especially for those that are lowly expressed. Only a handful of loci and developmental contexts, including *Shh* in vertebrate limb development^33–35^ and *Hbg/Hbb* in fetal and adult red erythropoiesis^36–38^ have been studied in great detail.

In this study, we provide a detailed account of postnatal, tissue-specific WNT gene expression regulation. We focus our efforts on the mammary gland, which has a complex and tightly regulated spatiotemporal *Wnt* gene expression landscape^39,40^. The primary function of the mammary gland is to produce and secrete milk for newborn offspring^41^. As such, it is critical for the survival of all mammalian species. In both mice and humans, WNT4 is important for mammary epithelial side-branching during the reproductive cycle and in early pregnancy. This is at least partly due to its activity in controlling the mammary stem cell population downstream of progesterone^42–45^. Here, we comprehensively map and dissect the mouse *Wnt4* and human *WNT4* cis-acting enhancer repertoire in the mammary gland and identify a conserved distal enhancer and regulatory chromatin hub in both mice and humans. We also categorize GRHL proteins as luminal transcription factors that control the recruitment of species-specific progesterone-dependent enhancers to regulate *Wnt4*/*WNT4* expression in mature luminal cells using two-step priming and transcriptional regulation at the chromatin level. Together, our data reveal how a subpopulation of mammary epithelial cells can convert a systemic endocrine signal into a local paracrine growth factor driven response. More broadly speaking, we present a model for how the dynamic interplay between physical chromatin looping and transcription factor binding can be directed to drive tissue-specific patterns of developmental gene expression.

## Results

### Identification of tissue-specific *Wnt4* enhancer candidates

The mammary gland is a dynamic tissue that undergoes most of its growth and differentiation postnatally. In the mouse, the mammary epithelium undergoes branching morphogenesis during puberty, invading the surrounding stroma to form an extensive, bilayered ductal network, consisting of basal and luminal cells (**Fig1A**). Throughout this time, *Wnt4* expression steadily increases^46^. The human breast shares many similarities with the mouse mammary gland at the molecular, cell and tissue level^47–50^. In both mice (**Fig1B, SuppFig1A-D**) and humans **(Fig1B, SuppFig1E**), we find *Wnt4*/*WNT4* expression to be largely restricted to a subset of luminal cells, predominantly composed of estrogen- and progesterone-responsive, mature luminal – or luminal hormone sensing – cells, as reported previously^39,42–45,51,52^. Indeed, WNT4 has mainly been reported to function downstream of progesterone^53,54^.

**Fig 1.**
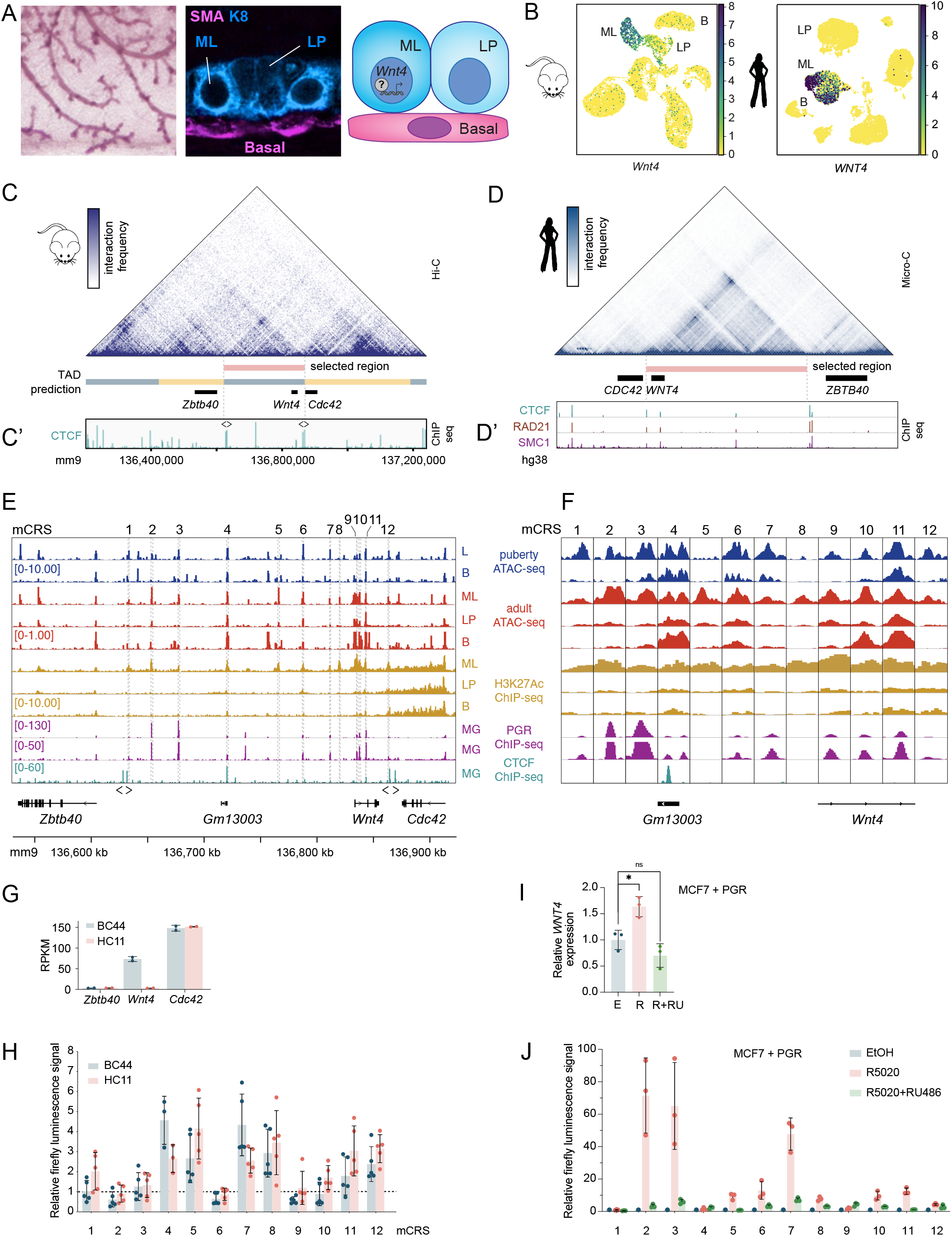
Identification of putative *Wnt4* cis-regulatory elements in the mouse mammary gland. A) Left: Wholemount, carmine-stained mouse mammary gland, showing part of the branched epithelial network. Middle: Confocal microscopy picture of the bilayered mammary epithelium, showing a close-up of luminal progenitor (LP, K8^low^), mature luminal (ML, K8^high^) and basal cells (SMA^+^). Right: *Wnt4* is expressed in progesterone-responsive ML cells. Secreted WNT4 protein signals to nearby basal cells. B) UMAP plots displaying scRNA-seq data from Tabula Muris Senis^136^ (left) and Tabula Sapiens^88^ (right), showing *Wnt4* (left) and *WNT4* (right) expression in ML cells in the mouse mammary gland (left) and human breast (right). Data visualization in CellxGeneVIP^137^. C) Hi-C plot depicting the murine *Wnt4* genomic region (mm9 chr4:136,200,000-137,250,000). Data from Rao et al.^58^, visualization performed in the 3D genome browser using the pipeline from Dixon et al.^138^ Cell type: CH12. TADs are predicted based on directionality index^56^. The selected *Wnt4* TAD is highlighted in pink (mm9 chr4:136,625,000-136,875,000). C’) CTCF ChIP-seq track displaying data from whole mouse mammary gland from Shin et al.^61^ Arrowheads indicate directional orientation of paired CTCF sites at the boundaries of the *Wnt4* TAD as determined manually using TRANSFAC^93,94,139^. D) Micro-C depicting the human *WNT4* genomic region (hg38 chr1:21,955,000-22,720,000). Data from human ES cells^140^, visualization performed in the 3D Genome Browser^138^. The selected *WNT4* TAD is highlighted in pink (hg38 coordinates chr1: 22,104,000-22,421,000). D’) CTCF^141,142^, RAD21^141,142^ and SMC1^143^ ChIP-seq tracks displaying data from human breast epithelial cells support the presence of the CTCF/COHESIN complex at the TAD boundaries. E-F) ATAC-seq and ChIP-seq tracks used to select mouse candidate regulatory sequences (mCRS) in the *Wnt4* TAD (mm9 coordinates of the depicted region chr4:136525000-136925000, data visualized in the IGV browser^144^). 12 mCRSs were selected based on higher ATAC-seq and/or H3K27ac ChIP-seq signal in ML cells compared to LP or B cells. mCRS numbering is in order of their genomic location. Coordinates of individual enhancers are provided in SuppTable1. Arrowheads highlight the orientation of CTCF motifs at the TAD boundaries. Puberty ATAC-seq dataset (blue) generated for this study. Adult ATAC-seq (red) and H3K27ac ChIP-seq (gold) data from adult virgin mice by Dravis et al.^145^, CTCF ChIP-seq data (mint green) from lactating mice by Shin et al.^61^, PGR ChIP-seq data (purple) from adult mice by Lain et al^146^ (displayed twice with different scales to highlight more prominent (top) and less prominent (bottom) peaks). F) is a close-up of the regions selected in E). L = Luminal cells, B= basal cells, ML = Mature luminal cells, LP = Luminal progenitor cells, MG = whole mammary gland, PGR = Progesterone receptor. G) Bar graph depicting mean relative expression of *Wnt4* and its neighbors, *Zbtb40* and *Cdc42*, measured by RNA-seq in BC44 (blue) and HC11 (red) mouse mammary epithelial cell lines (dataset generated for this study, n=2 replicates for each cell line). Datapoints show the n=2 individual values. RPKM = reads per kilobase per million mapped reads. H) Bar graph showing mean baseline enhancer activity of mCRS1-12 in BC44 (blue) and HC11 (red), measured in transient dual luciferase reporter assays. Empty vector control was set to 1. Datapoints: Individual values for n=4-5 independent biological experiments. Error bars: standard deviation (SD). I) Bar graph showing mean relative *WNT4* expression measured by qRT-PCR in MCF7 cells transiently transfected with *PGR* in the absence and presence of PR signaling. Reference gene: *YWHAZ*. EtOH-treated control (E, blue), R5020 stimulated (R, red) and R5020+RU486 treated (R+RU, green) samples were normalized to the average value of the control samples. Data points: Individual values for n=3 independent biological experiments. Error bars: SD. J) Bar graph showing mean relative progesterone-dependent enhancer activity of mCRS1-12, measured in transient dual luciferase reporter assays in MCF7 cells transiently transfected with *PGR*, following control (EtOH, blue), R5020 (red) or R5020+RU486 (green) treatment. Datapoints: Individual values for n=3 independent biological experiments. Error bars: SD.

To gain a better understanding of the molecular mechanisms responsible for the defined spatiotemporal control of WNT4 expression in the mammary gland, we sought to identify the responsible cis-regulatory sequences (CRS), hereafter also referred to as enhancers. Anticipating that this would require multimodal data integration, we carried out our initial search in the mouse, for the simple reason that compared to the human breast, the mouse mammary gland offers easier access to primary tissue of different developmental stages, and shows less donor-to donor heterogeneity due to the availability of inbred strains. Considering that both conserved and species-specific mechanisms might be at play, however, we cross-compared mouse *Wnt4* and human *WNT4* regulation throughout the study, both for discovery and validation purposes.

Interactions between enhancers and promoters primarily occur within large insulated regions of chromatin called topologically associating domains (TADs)^55–57^. To restrict our search area, we therefore first determined the boundaries of the *Wnt4*/*WNT4* TAD. Because no Hi-C data were available from primary mouse mammary gland or human breast tissue, and since TADs have been reported to be conserved across tissues and species^56,58^, we analyzed 91 publicly available mouse and human Hi-C datasets from different cell types and aligned their predicted *Wnt4*/*WNT4* TADs (**SuppFig1F**). Although the resolution and lengths of called TADs varied considerably across datasets, this approach allowed us to identify a minimally overlapping region of ∼250 kb in mice (**Fig1C**, mm9 coordinates chr4:136625000-136875000) and ∼310 kb in humans (**Fig1D**, hg38 coordinates chr1:22104249-22421001), which we defined as the minimal *Wnt4*/*WNT4* TAD for further analyses. As CTCF is commonly described to be present at TAD boundaries to facilitate loop formation^59,60^, we cross-referenced the *Wnt4* TAD with CTCF ChIP-seq data from whole mouse mammary gland^61^ (**Fig1C’**). This confirmed that the borders of the minimal *Wnt4* TAD are marked by CTCF binding to opposite facing pairs of CTCF sites, which is a signature of conserved TAD boundaries^62,63^. Of note, *Wnt4* is the only annotated protein coding gene in this TAD, with its neighboring genes *Zbtb40* and *Cdc42* being assigned to upstream and downstream TADs, respectively. The global organization of this minimal TAD, including flanking convergent CTCF and cohesion binding sites, is conserved in the human genome, although its orientation is inverted (**Fig1D’**).

We reasoned that lineage-specific *Wnt4* regulatory elements should be open and active^64– 67^ in mature luminal (ML) epithelial cells where *Wnt4* is predominantly expressed, rather than in luminal progenitor (LP) or basal (B) cells. Inspection of ATAC-seq and ChIP-seq data indicated 12 candidate sequences (mCRS1-12, full list in SuppTable1) for which H3K27 acetylation, a mark for active enhancers, was enriched in ML cells in adult mice (**Fig1E-F**, H3K27Ac ChIP-seq tracks). In most cases, chromatin accessibility also specifically increased in ML cells (**Fig1E-F**, adult ATAC-seq tracks), although some elements were open in all queried cell types (mCRS4, mCRS6, mCRS10 and mCRS11). Comparison of puberty and adult stages showed that the chromatin accessibility pattern present during puberty was maintained in the adult (**Fig1E-F**, ATAC-seq tracks). Most candidate enhancers lie upstream (mCRS1-8) of the *Wnt4* transcriptional start site (TSS), and three are intronic enhancers located within the *Wnt4* gene body (mCRS9, mCRS10, mCRS11). One enhancer (mCRS4) overlaps with the promoter and TSS of a long non-coding RNA (lncRNA), *Gm13003* (**Fig1E-F**).

### Candidate enhancers are both progesterone dependent and -independent

Of the 12 putative *Wnt4* regulatory elements only two (mCRS2 and mCRS3) display strong evidence of progesterone-receptor (PGR) binding in the mouse mammary gland *in vivo* (**Fig1E-F**, PGR ChIP-seq). To directly probe progesterone-dependent and –independent enhancer activity, we tested mCRS1-12 in either the absence or presence of active PGR signaling using luciferase reporter assays to determine the strength of cis-acting enhancer activity^68^ in multiple cell lines.

Baseline, progesterone-independent activity was measured in BC44 and HC11 mouse mammary epithelial cells, which are devoid of PGR expression^69,70^. Although *Wnt4* is expressed in both, the difference in mRNA expression levels is ∼25-fold (BC44: 73.4 RPKM, HC11: 3.0 RPKM, **Fig1G**). We hypothesized that this could reflect the availability of relevant transcription factors and, as such, these Wnt4^high^ (BC44) and Wnt4^low^ (HC11) conditions might constitute a more (BC44) or less (HC11) conducive environment for our candidate enhancers. No consistent difference in cis-regulatory activity was detected between the two cell lines, however (**Fig1H**). Some enhancers lacked baseline activity or even appeared to be repressive (e.g. mCRS2 and CRS6), while others gave modest (∼4-fold) activation (e.g. mCRS4 and mCRS7). Thus, variable enhancer activity exists in the absence of progesterone signaling.

Measuring progesterone signaling in cell lines is known to be challenging^53,71–75^. We recently optimized the experimental conditions to quantify functional PGR signaling activity in human breast cancer cell lines^76^, including MCF7, in which stimulation with the synthetic progestin R5020 reliably increased endogenous *WNT4* expression ∼1.5-2-fold following transient overexpression of PGR (**Fig1I**). Addition of RU486 (a competitive PGR inhibitor) abrogated this effect, demonstrating specificity. Using this same setup to screen our 12 candidate enhancers for PGR-dependent luciferase reporter activity, only mCRS2, 3 and 7 were clearly PGR-responsive relative to their baseline activity, showing >50-fold induction upon R5020 treatment (**Fig1J**). Taken together, we identified 12 new candidate enhancers in the *Wnt4* TAD. All elements are accessible and active in mature luminal mouse mammary epithelial cells (**Fig1E-F**), but only three are directly responsive to progesterone.

### mCRS4 physically interacts with the *Wnt4* promoter via chromatin looping

Although luciferase reporter assays allowed us to test baseline and progesterone-induced enhancer activity, they do not provide insight into how these sequences operate in a genomic context. As follow up, we therefore sought to functionally link individual elements to *Wnt4*. We first asked if any of the putative enhancer regions interacted with *Wnt4* via ‘enhancer-promoter’ looping^77–79^. To this end, we performed 4C chromatin conformation capture^80–82^, using the *Wnt4* promoter as a viewpoint (**Fig2A-B**). Because our 4C workflow required ∼10 million cells as input, we carried out this analysis using cell lines rather than primary cells. As expected, DNA regions within the *Wnt4* TAD displayed an overall higher interaction frequency than regions beyond the predicted TAD boundaries. Only one region, overlapping with mCRS4, showed statistically significant interactions with the *Wnt4* promoter above background (red shading, **Fig2A-B**) in both Wnt4^high^ (BC44) and Wnt4^low^ (HC11) cells.

**Fig 2.**
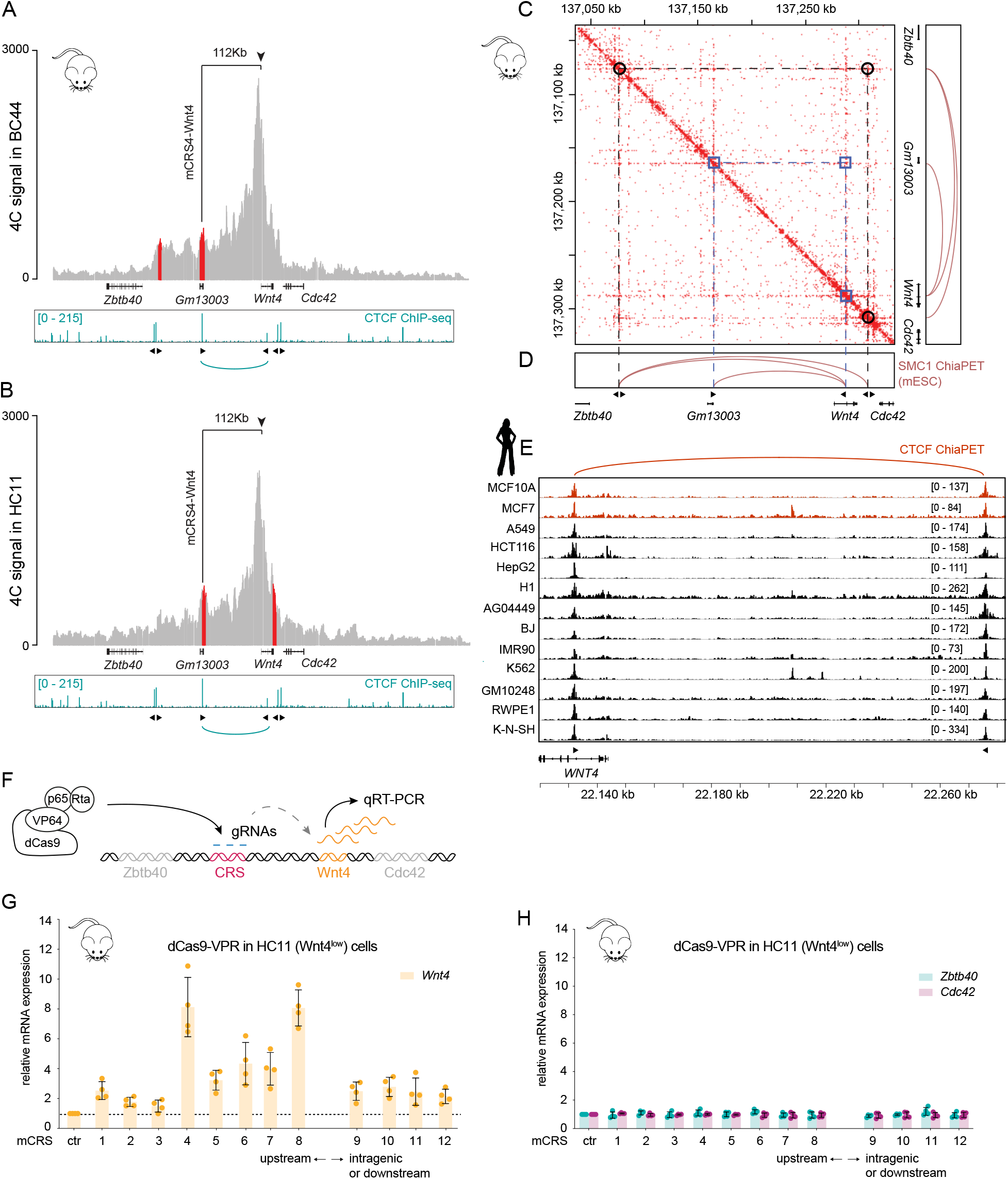
mCRS4 divides the *Wnt4* TAD into two separate loops. A-B) 4C plots showing a linear representation of the physical interactions between a viewpoint in the *Wnt4* promoter and other regions within the *Wnt4* TAD in BC44 (A) and HC11 (B) cells. For each cell line, cumulative analysis of n=3 biological experiments is shown. Regions with statistically significant interactions above the background model indicated in red. Data aligned to mm9 and analyzed in a window size of 400kb. The interaction between mCRS4 (112 kb upstream) and the *Wnt4* viewpoint (vertical arrowhead) is highlighted. Data visualized with PeakC^147^. Whole mammary gland CTCF ChIP seq^61^ tracks (same as in Fig1C) depicted below each graph for reference. Arrowheads below ChIP-seq tracks indicate orientation of CTCF sites. C-D) CTCF-ChiaPET heatmap (C) and SMC1 ChiaPET loops (D) confirm the presence of a CTCF/SMC1-mediated mCRS4-*Wnt4* loop in mouse embryonic stem cells. CTCF ChiaPET^141,142^ heatmap generated in JuiceBox^148^. SMC1 ChiaPET^149^ visualized in the 3D Genome Browser^138^. Black circles: boundaries of the *Wnt4* TAD. Purple squares: the mCRS4-*Wnt4* intra-TAD loop. Arrowheads: orientation of CTCF sites. E) Compilation of CTCF ChiaPET tracks from 13 human cell lines^141,142^ confirms structural conservation of a CTCF mediated intra-TAD loop (top, red). Linear visualization in the IGV browser^144^. Human breast (cancer) cell lines MCF7 and MCF10A highlighted in red. Data aligned to hg38. Arrowheads: orientation of CTCF sites. F) Cartoon depicting the strategy for guiding dCas9-VPR to a given enhancer (red) using specific gRNAs to test subsequent induction of endogenous *Wnt4* gene expression (yellow). If the enhancer/promoter contact is specific, the transcriptional activation signal should not be transferred to neighboring genes (grey). G-H) Bar graphs showing specific induction of *Wnt4* (G, yellow) but not its neighboring genes (H) *Zbtb40* (mint green) or *Cdc42* (purple) measured by qRT-PCR upon guiding dCas9-VPR to individual mCRSs in HC11 (Wnt4^low^) cells. mCRSs are ordered along the x-axis according to their genomic location. Ctr = control without gRNAs, set to 1. Reference gene: *Ctbp1*^46^. Bars: relative mean expression. Datapoints: Individual values for n=4 biological replicates. Errors bars: SD.

mCRS4 is located more than 100 kb upstream of *Wnt4*. Upon closer inspection, we noticed that this element is bound by CTCF in the mouse mammary gland (**Fig1F**), suggesting direct physical looping between mCRS4 and *Wnt4*. Indeed, analysis of CTCF (**Fig2C**) and SMC1 (**Fig2D**) ChiaPET data from mouse embryonic stem cells confirmed that a stable, CTCF and Cohesin mediated 112 kb intra-TAD loop involving mCRS4 can form inside the larger *Wnt4* TAD. Given that this loop is present in each of the settings we tested, we conclude that this is a stable loop that forms irrespective of *Wnt4* expression levels. This structural chromatin configuration is conserved in humans, as demonstrated by CTCF, RAD21 and SMC1 ChIP-seq (**Fig1D’**), as well as by CTCF ChiaPET data obtained from human cell lines of multiple different origins (**Fig2E, SuppFig3A**).

To further validate that one or more of our candidate enhancers are in direct contact and communication with *Wnt4*, we performed transient CRISPR activation assays (CRISPRa) using a nuclease-dead Cas9 fused to a strong transcriptional activator (dCas9-VPR)^83,84^. We designed tiling sets of sgRNAs to direct dCas9-VPR to each mCRS (∼1 sgRNA per 150bp) and measured the change in expression of *Wnt4* by qRT-PCR (**Fig2F**). We hypothesized that using this setup, only enhancer sequences in close 3D proximity to the *Wnt4* promoter should be capable of activating endogenous *Wnt4* expression. As controls we measured expression of the *Wnt4* neighboring genes, *Cdc42* and *Zbtb40*, which should not be affected by dCas9-VPR targeting of *Wnt4*-specific enhancers.

In BC44 (*Wnt4*^*high*^) cells, the effect of dCas9-VPR mediated activation of *Wnt4* and neighboring gene expression was minor (**SuppFig2A-B**). This is in line with reports in the literature that describe a negative correlation between endogenous gene expression levels and dCas9-VPR inducibility^83,84^. In HC11 (*Wnt4*^*low*^) cells, however, recruitment of dCas9-VPR to 10 out of 12 mCRSs induced endogenous *Wnt4* expression >2-fold (**Fig2G**). The distal enhancer mCRS4 increased endogenous *Wnt4* gene expression 8.1-fold (even though it is located ±100 kb upstream of the *Wnt4* TSS), comparable to the induction achieved by recruiting dCas9-VPR to mCRS8 (which lies in much closer proximity to the *Wnt4* TSS, ±10 kb upstream, 8.0-fold mean increase in endogenous *Wnt4* expression). In all cases, expression of *Cdc42* and *Zbtb40* was unaffected (**Fig2H**). Summarizing, the link between mCRS4 and *Wnt4* is specific and exclusive, although other elements also contribute to driving *Wnt4* expression. The PGR-dependent candidate enhancers mCRS2 and mCRS3 were not capable of inducing endogenous *Wnt4* expression using our CRISPRa setup (**Fig2G, SuppFig2A**), suggesting that, at least in the absence of active progesterone signaling, these sequences do not come in close physical proximity to the *Wnt4* promoter.

### Identification of a conserved regulatory chromatin hub in mice and humans

Based on our combined findings thus far, we aimed to further dissect the enhancer hierarchy for both mouse and human. To validate that mCRS4 and *Wnt4* also form a stable loop *in vivo*, we re-analyzed previously published mouse mammary single cell ATAC-seq data (scATAC-seq)^85^ using the Cicero algorithm^86^, which predicts putative cis-regulatory interactions based on single-cell chromatin co-accessibility of distal DNA elements and their presumed target gene. We identified two prominent interactions with the *Wnt4* promoter: the distal enhancer mCRS4 and the intragenic enhancer mCRS9 (**Fig3A**). Of these, mCRS4 had the highest score and confidence, thus confirming that the chromatin loop identified in cell lines (**Fig2A-B**) is also present *in vivo*.

**Fig 3.**
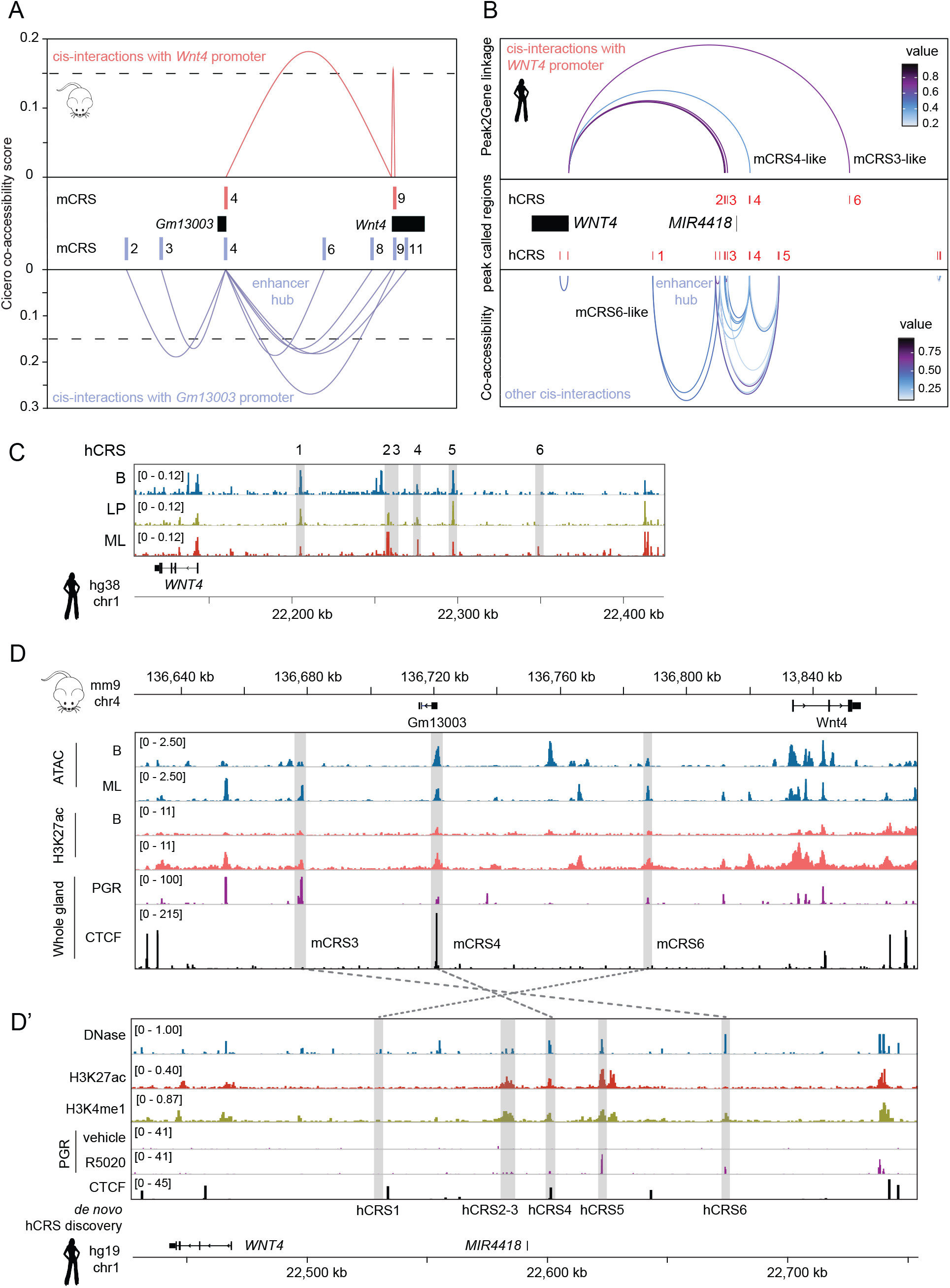
Conserved and species-specific enhancers converge on a regulatory chromatin hub to regulate mouse *Wnt4* and human *WNT4* expression. A) Cicero co-accessibility plot highlighting inferred cis-regulatory element and chromatin interactions in the mouse mammary gland *in vivo*. Only mCRS4 and mCRS9 are predicted to interact with the *Wnt4* promoter. Multiple other mCRSs are predicted to interact with mCRS4. Data from Chung et al.^85^. Aligned to mm10. B) Peak2Gene linkage and co-accessibility plot showing cis-regulatory elements that are linked to *WNT4* in the human breast *in vivo*. hCRS4 is part of a larger enhancer hub. Data from Zhang et al.^87^ were analyzed and visualized in ArchR^150^. Aligned to hg38. D) Pseudobulk scATAC-seq tracks from different human breast epithelial cell populations visualized linearly in the *WNT4* TAD. LP = luminal progenitor, ML = mature luminal. B = basal. Grey columns highlight genomic regions of conserved hCRSs identified in (B). Data from Zhang et al.^87^, visualized in the IGV browser^144^. Aligned to hg38. D/D’) Alignment of mouse mammary gland ATAC-seq and ChIP-seq (D) and human breast DNAse-seq and ChIP-seq (D’) tracks, illustrating conservation of epigenetic features and cis-regulatory elements. Depicted human breast data are from MCF-7 cells (DNase-seq and CTCF ChIP-seq from ENCODE^141,142^, H3K27ac and H3K4me1 from Gala et al.^151^, PGR from Mohammed et al.^73,152^). Mouse ATAC-seq and H3K27ac ChIP-seq from Dravis et al.^145^, whole mammary gland CTCF ChIPseq from Shin et al.^61^ and whole mammary gland PGR ChIP-seq from Lain et al.^146^. Data visualized in the IGV browser^144^. Aligned to mm9 and hg19. Note reverse orientation of the mouse *Wnt4* and human *WNT4* locus. Genomic locations of the conserved mCRS3,4,6 are highlighted in dark grey in (D). Genomic locations of the human cis-regulatory elements (hCRS1-6) newly identified in (B) are also highlighted in dark grey (D’). Conserved sequences (as identified by BLAST^153^) mCRS3, mCRS4 and mCRS6, correspond to hCRS6, hCRS4 and hCRS1, respectively (dashed lines). ML = mature luminal. B = basal.

If the aforementioned murine lncRNA *Gm13003* (**Fig1E-F**) is transcribed from the active mCRS4 enhancer due to close physical proximity to the *Wnt4* promoter, one would expect *Gm13003* to closely follow *Wnt4* gene expression. Indeed, *Gm13003* transcript levels are higher in BC44 than in HC11 cells, similar to the *Wnt4* expression pattern (**SuppFig2C**). *Gm13003* and *Wnt4* expression also correlate *in vivo*, as shown by qRT-PCR analysis of FACS sorted primary mammary epithelial cell populations (**SuppFig2D**). Interestingly, when the *Gm13003* promoter is used as a viewpoint (or anchor) for Cicero analysis, mCRS4 is predicted to not only interact with the *Wnt4* promoter, but also with 6 other candidate enhancers that we identified in our initial screen (**Fig3A**). We therefore propose that mCRS4 not only forms a stable, physical loop with *Wnt4*, but also lies at the heart of a regulatory chromatin hub that attracts and facilitates close interactions with other upstream (PGR-responsive mCRS2 and mCRS3) and downstream cis-regulatory elements (mCRS6, mCRS8, mCRS9 and mCRS11).

Given the presence of a conserved CTCF-mediated intra-TAD loop in humans (**Fig1D’** and **Fig2E**), we asked if a similar functional conformation might exist in the human genome. To this end, we first integrated human scATAC-seq^87^ and scRNA-seq data^88^ for unbiased, *de novo* discovery of cis-regulatory elements in the minimal human *WNT4* TAD (**Fig3B**). From this, we inferred *in vivo* DNA interactions as in **Fig3A**. Similar to the mouse, a distal chromatin hub upstream of *WNT4* is predicted to pull in both upstream and downstream enhancers. Note that the human hub spreads out over a larger region and extends beyond a single individual enhancer element (**Fig3B**).

We identified 6 putative human *WNT4* cis-regulatory elements (labeled hCRS1-6 in **Fig3B-D**, full list in SuppTable1) for further follow up based on their combined DNA accessibility, histone modification and transcription factor binding features in primary tissue (**Fig3C**) and cell lines (**Fig3D’**). Four of these elements (hCRS2-4 and hCRS6) show enriched chromatin accessibility in ML primary human breast epithelial cells (**Fig3C**). Three elements forming the distal chromatin hub (hCRS2-4, ±20 kb) are predicted to directly loop to the human *WNT4* promoter (**Fig3B**). Of these, only hCRS4 shows evidence of CTCF binding (**Fig3D’** and **SuppFig3A**), suggesting that it plays a critical role in forming and maintaining the intra-TAD loop identified earlier (**Fig1D’** and **Fig2E**).

Of note, hCRS4 shows sequence conservation with mCRS4 beyond the presence of a CTCF site. Two other elements (hCRS1 and hCRS6) are also conserved at the DNA sequence level (hCRS1 with the PGR-independent mCRS6 and hCRS6 with the PGR-dependent mCRS3 (**Fig3D-D’**). Finally, hCRS5 is not conserved with any of the mouse enhancer sequences at the DNA sequence level but does show evidence of PGR binding in MCF7 cells (**Fig3D’**) and could thus function as a species-specific progesterone-responsive enhancer, as is the case for mCRS2 in the mouse. Indeed, we can confirm baseline (**SuppFig3B**) and/or PGR-dependent (**SuppFig3C**) enhancer activity for each of the six human enhancer sequences, with the conserved hCRS6 (mCRS3-like) showing the highest activation in response to R5020 treatment (**SuppFig3C**). Taken together, comprehensive dissection of the mouse *Wnt4* and human *WNT4* TAD allowed us to discover both conserved (mCRS3/hCRS6, mCRS4/hCRS4 and mCRS6/hCRS1) and species-specific enhancers (**SuppTable 1**) that are predicted to control *Wnt4/WNT4* expression in mature luminal epithelial cells in the mouse mammary gland and the human breast in a PGR-dependent and PGR-independent manner.

### GRHL proteins are luminal lineage factors that regulate *WNT4* expression through the conserved enhancer mCRS4/hCRS4

Based on our PGR binding (**Fig3D-D’**) and PGR signaling analyses so far (**Fig1J** and **SuppFig3C**), progesterone is unlikely to be the sole driver of *WNT4* expression in either the mouse mammary gland or the human breast. Therefore, we used an unbiased approach to not only discover novel *WNT4* regulatory factors but also link these factors to the individual cis-regulatory elements we identified in this study. We reasoned that a candidate transcription factor should meet two criteria. First, the gene encoding the transcription factor should be both accessible and expressed in cells that express *WNT4*, i.e. ML cells (**Fig4A**, top). Second, regions harboring the relevant transcription binding motif (including some of our enhancer sequences) should themselves be open and enriched in this cell population, with functional evidence of DNA binding by the candidate transcription factor at its cognate motif (**Fig4A**, bottom). To assess each of these criteria, we reanalyzed and integrated available scATAC-seq, scRNA-seq and ChIP-seq datasets in a stepwise manner (**Fig4B, SuppFig4**).

**Fig 4.**
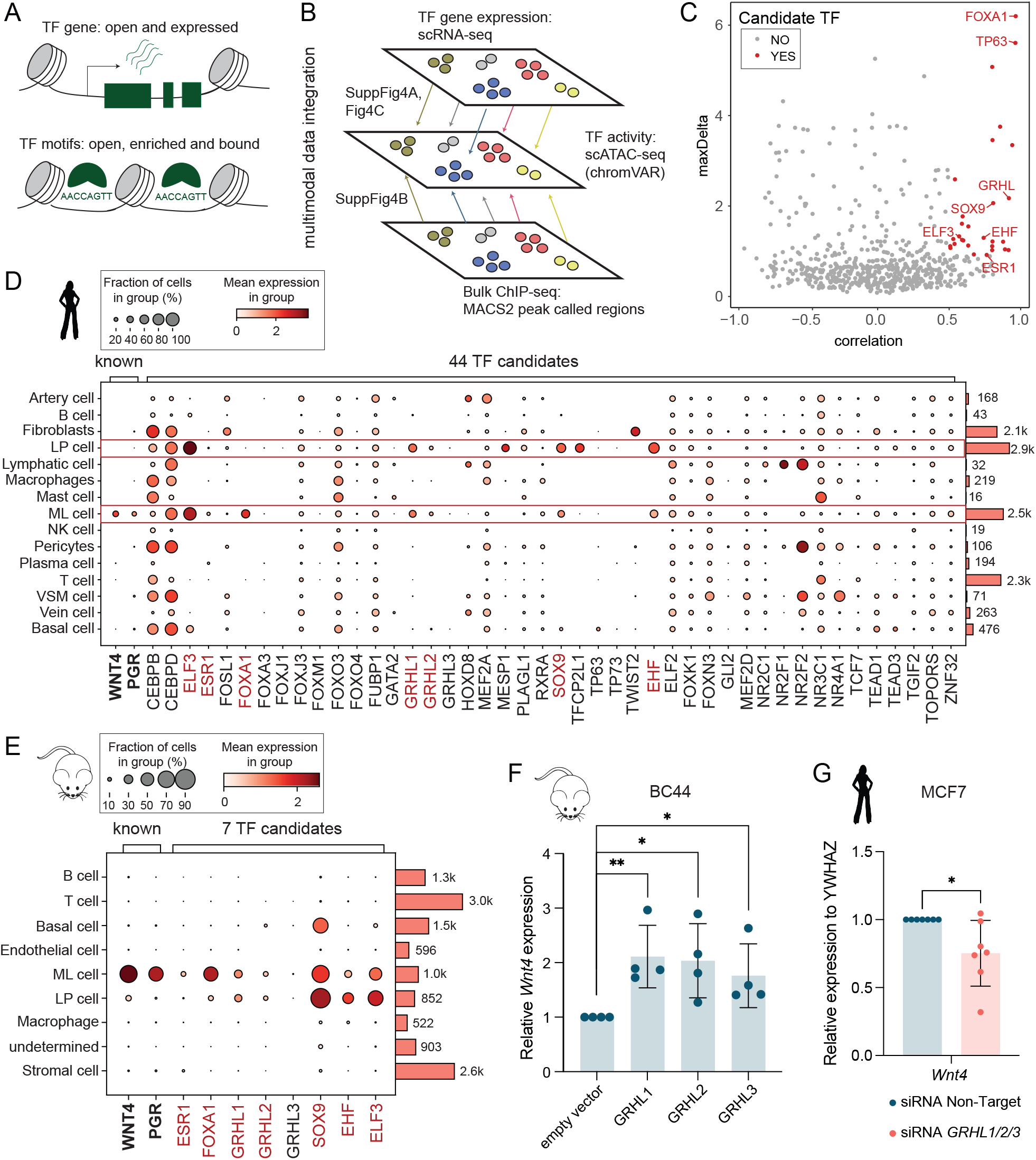
Identification of GRHL transcription factors as novel *WNT4* regulators. A) Cartoon depicting criteria to qualify as candidate regulatory transcription factor (TF). TF gene expression, motif accessibility and DNA binding must be enriched in one cell cluster compared to others (here: basal, luminal progenitor or mature luminal breast epithelial cell clusters). B) Cartoon depicting workflow for multimodal integration of scRNA-seq, scATAC-seq and ChIP-seq data in UMAP space using ArchR^150^ to predict and confirm preferential expression and binding of lineage-specific candidate regulatory TFs in different cell clusters. (C) Scatterplot depicting results from CHROMVAR^89^ analysis. Binding motif enrichment was performed on primary human breast scATAC-seq data to predict TF activity in distinct human breast cell populations (maxDelta: difference between clusters). This was correlated to primary human breast scRNA-seq data to identify TFs whose predicted activity and gene expression correlate (correlation score). Candidate TFs that pass the threshold (correlation >0.5, p-adj. value <0.01, maxDelta deviation z-score in the top quartile) are highlighted in red and include known lineage factors such as FOXA1, TP63 and ESR1. Depicted human scATAC-seq data from Zhang et al.^87^ and scRNA-seq data from Bhat-Nkshatri et al.^90^. D-E) Bubble plot depicting candidate TF expression in different human breast (D, scRNA-seq data from Tabula Sapiens^88^) and mouse mammary gland (E, scRNA-seq from Tabula Muris Senis^154^) cell populations visualized in CellxGeneVIP^137^. D: 44 candidate TFs identified in (C). E: 7 shortlisted TFs. Columns: Individual genes. *WNT4/Wnt4* and *PGR/Pgr* included for reference. Rows: Different cell populations (based on scRNA-seq clustering). Bubble size: fraction of cells in each cluster expressing the TF. Color: mean expression within this fraction. Bars on the right indicate the number of cells within each cluster. F) Bar graph depicting mean relative *Wnt4* gene expression measured by qRT-PCR after transient overexpression of GRHL1, GRHL2 or GRHL3 in serum starved mouse mammary epithelial BC44 cells. Reference gene: *Rpl13a*. Data were normalized to an empty vector control for each biological experiment. Datapoints: Individual values for n=4 biological repeats. Asterisks: statistical significance (* = p < 0.05, ** = p <0.01). Error bars: SD. G) Bar graph depicting mean relative *WNT4* gene expression measured by qRT-PCR following transient siRNA knockdown of *GRHL1, 2 and 3* in human breast epithelial MCF7 cells. Reference gene: *YWHAZ*. Data were normalized to the non-targeting siRNA control for each biological experiment. Datapoints: Individual values for n=7 biological repeats. Asterisks: statistical significance (* = p < 0.05). Error bars: SD.

First, we used chromVAR^89^ to analyze scATAC seq data^87^ in order to identify transcription factor binding motifs of which the accessibility is enriched in different human breast cell populations. We correlated this motif accessibility enrichment analysis to transcription factor gene expression enrichment by integrating two different human breast scRNA-seq datasets^88,90^ (**SuppFig4A**). From this, we identified candidate transcription factors whose expression showed a positive correlation to the accessibility of their corresponding motif (**Fig4C**). This initial list contained 44 transcription factors and includes *TP63, ESR1* and *FOXA1*, all of which are known regulators of cell identity in the mammary gland (**Fig4D**).

Cross-referencing to human breast scRNA-seq data from the Tabula Sapiens Consortium^88^ (**SuppFig5**) allowed us to select transcription factors whose expression and activity was restricted to the luminal lineage, and preferably enriched in ML cells – thus matching the known *WNT4* and *PGR* pattern. This resulted in a final list of 7 candidate genes, including known luminal-specific transcription factors (*ESR1* and *FOXA1*), but also less well studied candidate regulators (*GRHL1, GRHL2, SOX9, EHF* and *ELF3*). Whereas *PGR, ESR1* and *FOXA1* expression is largely restricted to ML cells, *GRHL1, GRHL2, SOX9, EHF* and *ELF3* are more broadly expressed across the LP and ML cell populations (**Fig4D, SuppFig5)**. This expression pattern is conserved in the mouse mammary gland (**Fig4E**), indicating a potentially conserved role in mammary gland homeostasis. By analyzing pooled ChIP-seq data for the human breast, we found GRHL1 and GRHL2 DNA binding to also be enriched in ML and, to a lesser extent, LP cells (**SuppFig4B**). Based on these combined findings, we focused our further efforts on GRHL.

Both the mouse and the human genome encode three GRHL paralogues, with known functions in regulating epithelial morphogenesis and differentiation^91,92^. Transient overexpression of either GRHL1, GRHL2 or GRHL3 is sufficient to induce endogenous *Wnt4* expression in serum starved BC44 mouse mammary epithelial cells (**Fig4F**). Simultaneous siRNA knockdown of endogenous *GRHL1, GRHL2* and *GRHL3* expression in MCF7 cells resulted in an average 30% reduction of endogenous *WNT4* expression (**Fig4G**). Considering that endogenous *GRHL3* is expressed at very low levels (**Fig4D-E**), *GRHL1* and *GRHL2* are most likely to play a role under physiological conditions.

To dissect the regulatory mechanisms involved, we circled back to our individual enhancers and had a closer look at the DNA conservation between mCRS4 and hCRS4, the CTCF binding element that plays a central role in chromatin organization in the *Wnt4*/*WNT4* TAD (**Fig2A-E, Fig3A-B**). PhastCons and BLAST alignment to the human genome revealed two smaller conserved regions (**Fig5A-B, SuppFile1**). Both are open and accessible in mouse mammary gland and human breast epithelial cells (mouse ATAC-seq, **Fig5A** and human DNAseI-seq, **Fig5B**). Region 1 is responsible for the previously observed CTCF binding (**Fig1C’-D’**) and harbors a 25 bp element conserved in placental mammals with a CTCFL motif (Jaspar matrix ID MA1102.2) in mouse mCRS4 and a larger CTCF motif (Jaspar matrix ID MA1929.2) in human hCRS4 (**Fig5A-B, SuppFile1**). Region 2 is slightly larger and, in the mouse, falls within the *Gm13003* promoter (**Fig5A**). TRANSFAC analysis^93,94^ revealed the presence of a conserved GRHL binding motif at this location (**Fig5A-B, SuppFile1**). Indeed, GRHL2 binds to this region in both mouse (**Fig5C**) and human (**Fig5D**) cells and tissues.

**Fig 5.**
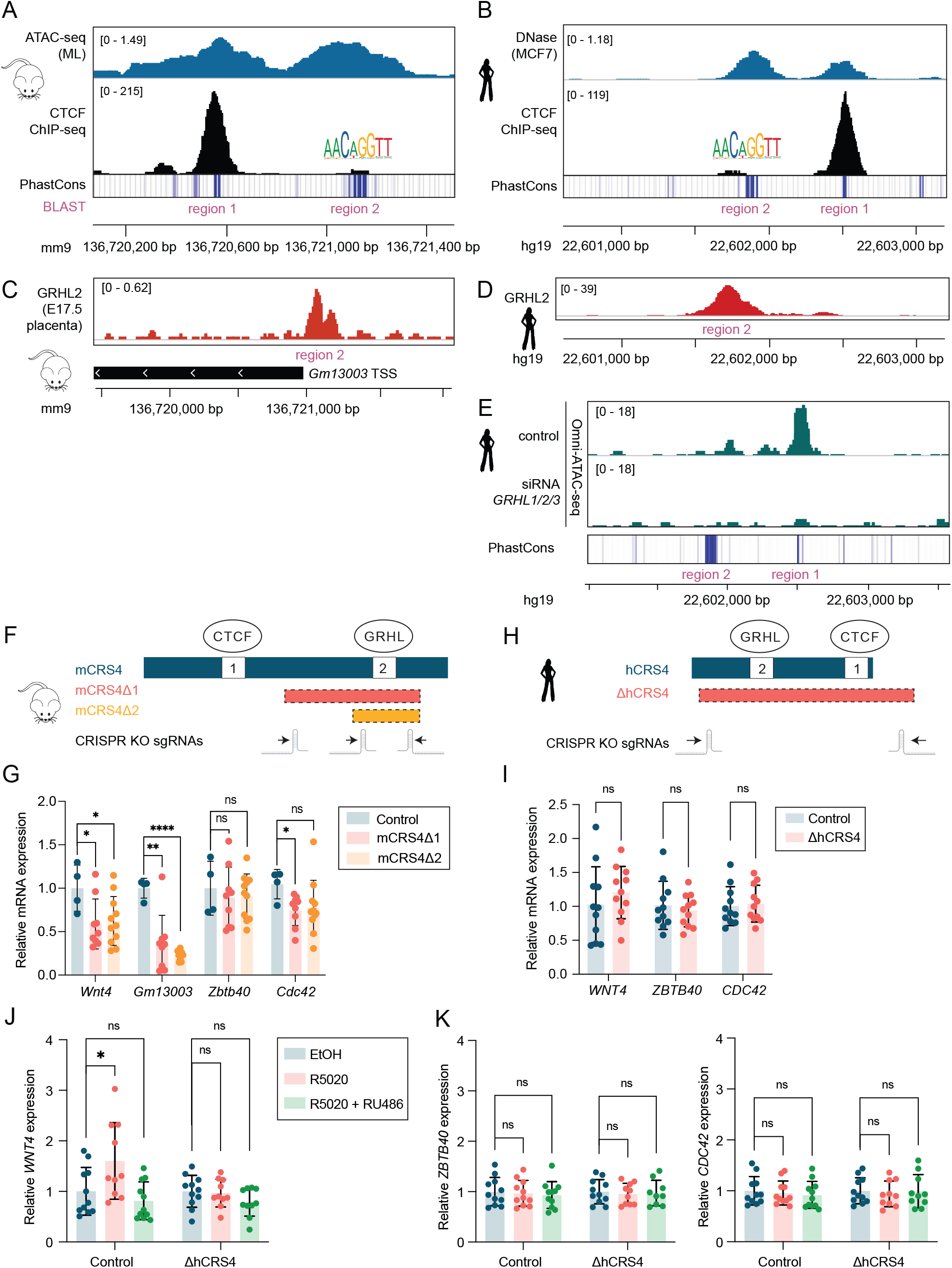
Context dependent changes in *Wnt4/WNT4* expression upon deletion of mCRS4/hCRS4. A-B) Conserved chromatin accessibility (ATAC-seq, regions 1 and 2), CTCF binding (ChIP-seq, region 1) and presence of a GRHL motif (region 2) in mouse mCRS4 (A) and human hCRS4 (B, inverted orientation compared to mouse) genomic region. Detailed sequence conservation of regions 1 and 2 identified by BLAST^153^ provided in SuppFile1. A) ATAC-seq (ML cells, data from Dravis et al.^145^), CTCF ChIP-seq (whole mammary gland, data from Shin et al.^61^) and PhastCons^155^ (data from UCSC) tracks. *(legend continues on next page)* B) DNAse-seq and CTCF ChIP-seq (MCF7 cells, data from ENCODE^141,142^ and Gala et al.^151^) and PhastCons^155^ (data from UCSC) tracks. C-D) GRHL2 ChIP-seq shows GRHL2 binding to conserved region 2 in mCRS4 (C, (E17.5 mouse placenta, data from Walentin et al.^156^, *Gm13003* TSS depicted for orientation) and hCRS4 (D, MCF7 cells, data from Cocce et al.^157^). E) Close-up of hCRS4, displaying Omni-ATAC-seq tracks from MCF7 cells (data from Jacobs et al.^95^) before (top) or after (bottom) *GRHL1/2/3* siRNA knockdown. Conservated regions 1 and 2 indicated for reference. F) CRISPR/Cas9 mediated knockout strategy for mCRS4 in BC44 cells, resulting in two targeted deletions: a 536 bp deletion removing both conserved region 2 and the *Gm13003* promoter plus TSS (mCRS4Δ1) and a 170 bp deletion comprising only conserved region 2 and the *Gm13003* promoter (mCRS4Δ2). Orientation (arrows) and location (hairpins) of the CRISPR gRNAs is indicated. G) Bar graph depicting the impact of the mCRS4 deletions from (F) on the relative normalized mean expression of *Wnt4, Gm13003* and the neighboring genes *Zbtb40* and *Cdc42* as measured by qRT-PCR. Data show combined results from n=2 CRISPR control clones (n=4 individual datapoints total), n=3 CRISPR mCRS4Δ1 clones (n=9 individual datapoints total) and n=4 CRSIPR mCRS4Δ2 clones (n=11 datapoints total). Reference genes: *Rpl13a, Cbtbp1* and *Prdx1*^46,158^. Asterisks: statistical significance (* = p < 0.05, ** = p <0.01, **** = p < 0.001, ns = not statistically significant). H) CRISPR/Cas9 mediated knockout strategy for hCRS4 in MCF7 cells. A targeted knockout (ΔhCRS4) encompassing both conserved region 1 (CTCF binding) and 2 (GRHL2 binding) was generated. Orientation (arrows) and location (hairpins) of the CRISPR gRNAs is indicated. I) Bar graph displaying relative mean baseline gene expression of *WNT4, CDC42* and *ZBTB40* in MCF7 ΔhCRS4 clones (n = 2) compared to control clones (n = 2) as measured by qRT-PCR. Reference gene: *YWHAZ*. All values depicted relative to the average expression in the control clones. Datapoints: Combined individual values for n=5 biological repeats per clone). Asterisks: Statistical significance (* = p < 0.05, ** = p <0.01, **** = p < 0.001, ns = not statistically significant). J) Bar graph of qRT-PCR data, demonstrating loss of *WNT4* induction in response to PGR signaling (R5020 treatment, red) in MCF7 ΔhCRS4 clones compared to control clones (same clones as in I). Treatment with R5020 and RU486 (green) shows specificity of the response. Reference gene: *YWHAZ*. All values depicted as mean fold change to the average value of the control (EtOH-treated, blue) samples. Datapoints: Individual values for n=5 biological repeats per clone. Asterisks: Statistical significance (* = p < 0.05, ** = p <0.01, **** = p < 0.001, ns = not statistically significant). K) Same as (J) but showing that expression of the neighboring genes *ZBTB40* (left) or *CDC42* (right) is unaffected by either R5020 treatment or hCRS4 loss.

Given the proposed role of GRHL proteins as pioneer transcription factors^91^, which are capable of opening closed chromatin, we asked if loss of GRHL function changed chromatin accessibility of the mCRS4/hCRS4 enhancer. Indeed, transient knockdown of *GRHL1, GRHL2* and *GRHL3* reduces chromatin accessibility in human MCF7 breast cancer cells^95^, specifically at the conserved CTCF site in region 1 immediately adjacent to the GRHL2 binding site (**Fig5E**). Thus, we conclude that GRHL proteins are both necessary and sufficient to regulate endogenous expression of mouse and human *Wnt4/WNT4* (**Fig4F-G**). This activity is, at least in part, due to their capacity to directly maintain an open chromatin state at mCRS4/hCRS4 (**Fig5A-E**).

To test if the mCRS4 element itself is required for baseline, progesterone-independent *Wnt4* expression, we used CRISPR/Cas9 gene editing to knock out mCRS4 in BC44 (*Wnt4*^*high*^) cells. We succeeded in generating a 536 bp deletion that removed both the *Gm13003* TSS and promoter (mCRS4Δ1), and a 170 bp deletion comprising only the *Gm13003* promoter (mCRS4Δ2, **Fig5F**). Even though the CTCF site was left intact, both mCRS4Δ1 and mCRS4Δ2 caused a statistically significant reduction in the expression of *Gm13003* and *Wnt4*, but not the neighboring genes *Zbtb40* and, to a lesser extent, *Cdc42* (**Fig5G**). The 170 bp deletion comprising the conserved GRHL binding site is sufficient to reduce *Gm13003* expression ∼4-fold (75% reduction) and *Wnt4* expression almost 2-fold (40% reduction).

To validate the importance of hCRS4 for *WNT4* expression in the human breast, we generated a 1400 bp CRISPR knockout of the hCRS4 element in MCF7 cells. This knockout encompasses both conserved regions, thus abrogating GRHL2 as well as CTCF binding (**Fig5H**). Next, we measured the *WNT4* expression levels in two independent CRISPR knockout and two independent control wildtype MCF7 clones. While deletion of hCRS4 did not affect basal *WNT4* expression (**Fig5I**), knockout clones were severely impaired in their capacity to induce endogenous *WNT4* expression in response to progesterone signaling compared to wildtype control clones, which showed a modest yet consistent 1.5-2-fold increase in *WNT4* expression upon R5020 treatment (**Fig5J**). As expected, expression of the neighboring genes *ZBTB40* and *CDC42*, which are not PGR-responsive, remained unaltered in both the absence and presence of R5020 treatment (**Fig5K**).

Although it is challenging to measure the *in vivo* effects of enhancer deletions^96^, we also knocked out the mCRS4 element in mice, generating two independent CRISPR knock-out lines on an FVB/N background with a 1351 bp and a 1352 bp deletion allele (**Fig6A**). Homozygous *Wnt4*Δ*mCRS4* mice were viable and fertile, were born in the expected Mendelian ratios (**SuppTable2**) and lived without any overt phenotype when followed up to 12 months of age. Homozygous *Wnt4*Δ*mCRS4* females showed no apparent delay in mammary gland outgrowth during puberty (**Fig6B**,**C**), nor a a consistent measurable change in *Wnt4* gene expression during this time (**Fig6D-F**). Finally, homozygous *Wnt4*Δ*mCRS4* females of mixed ages (16-25 weeks) showed no apparent delay in alveolar development at day 8.5 of pregnancy, when *Wnt4* expression drives alveolar side branching^42^ (**Fig6G**). Consistent with a lack of phenotype, we did not measure a reduction in *Wnt4* expression in homozygous mice compared to wildtypes at day 8.5 of pregnancy, when it normally peaks^97^ (**Fig6H**).

**Fig 6.**
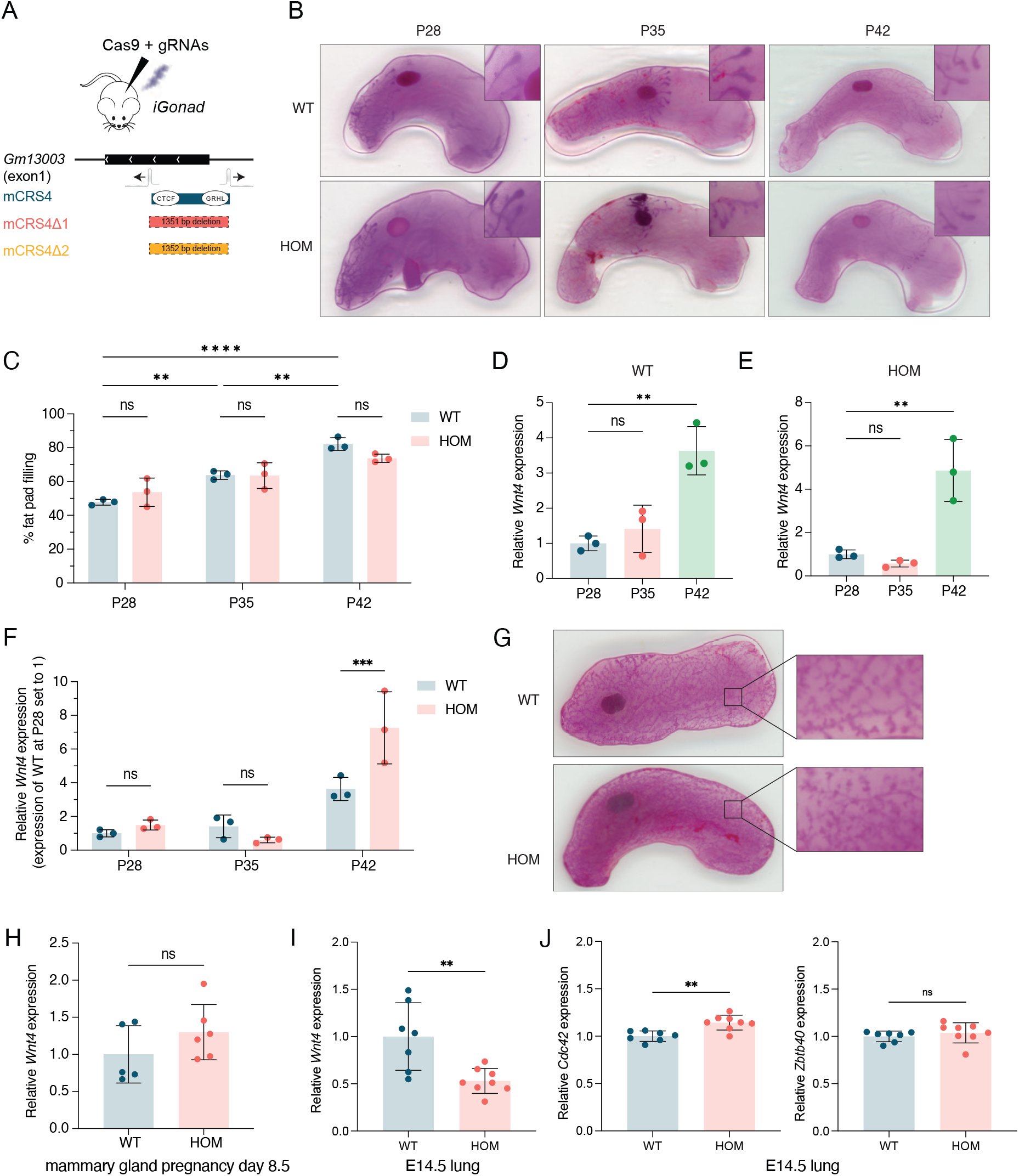
Deletion of mCRS4 mildly affects *Wnt4* gene expression *in vivo*. A) CRISPR/Cas9 mediated strategy for the mCRS4 *in vivo* knockout allele. Two independent knockout lines were generated, containing a 1351 bp (mCRS4Δ1) and a 1352 bp (mCRS4Δ2) deletion. Both remove the CTCF and GRHL binding sites including the *Gm13003* promoter and TSS. Orientation (arrows) and location (hairpins) of the CRISPR gRNAs is indicated. Panels B-I contain merged data from both alleles. B) Representative images of wholemount carmine alum stainings showing ductal outgrowth of pubertal (P28, P35 and P42) wildtype (WT, top) and homozygous ΔmCRS4 (HOM) mammary glands. Close-ups of terminal end buds at the leading edge of the ductal outgrowth displayed in the upper right corners. C) Bar graph with quantification of the data shown in (B), calculated as percentage fat pad filling at P28, P35 and P42 for wildtype (WT, blue) and homozygous ΔmCRS4 (HOM, red) mammary glands. For each developmental stage, n=3 WT and HOM mice were compared. Asterisks: statistical significance (* = p < 0.05, ** = p <0.01, **** = p < 0.001, ns = not statistically significant). D-F) Bar graphs of qRT-PCR data, depicting relative expression of *Wnt4* in wildtype (WT) and homozygous ΔmCRS4 (HOM) whole mammary glands obtained from mice at pubertal timepoints P28, P35 and P42 (n=3, contralateral glands of samples analyzed in B C). Reference gene: *β-actin*. D-E: Data for WT (D) and HOM (E) mice, normalized to mean *Wnt4* expression at P28 for each genotype. F: Joint data from D-E, showing WT (blue) and HOM (red) normalized to mean *Wnt4* expression in WT mice at P28. Asterisks: statistical significance (* = p < 0.05, ** = p <0.01, **** = p < 0.001, ns = not statistically significant). G) Representative images of wholemount carmine alum stainings showing ductal side branching in wildtype (WT, top) and homozygous ΔmCRS4 (HOM) mammary glands from 17-week old WT and 23-week old HOM mice at day 8.5 of pregnancy. H) Bar graph with qRT-PCR data, showing mean relative *Wnt4* gene expression levels in wildtype (WT, blue) and homozygous ΔmCRS4 (HOM) whole mammary glands from 16-25 week old mice at day 8.5 of pregnancy. Reference gene: β*-actin*. Datapoints: values for n=5-6 individual mice depicted as fold change compared to the average *Wnt4* expression in 8.5 day pregnant WT mammary glands. Asterisks: statistical significance (* = p < 0.05, ** = p <0.01, **** = p < 0.001, ns = not statistically significant). I-J) Bar graphs with qRT-PCR data, depicting mean relative expression of *Wnt4* (I) or its neighboring genes *Zbtb40* and *Cdc42* (J) in wildtype (WT, blue) and homozygous ΔmCRS4 (HOM, red) E14.5 lungs. Reference gene: *β-actin*. Datapoints: values for n=7 WT and n=8 HOM littermates from n=2 independent timed mating experiments are depicted as mean fold change relative to the average expression of each gene in WT embryonic lungs. Asterisks: statistical significance (* = p < 0.05, ** = p <0.01, **** = p < 0.001, ns = not statistically significant).

Given the known role of *Wnt4* in other developmental processes^98–102^, we also measured the effects of mCRS4 loss on *Wnt4* expression in other tissues. In homozygous *Wnt4*Δ*mCRS4* embryonic lungs, we were able to detect an approximately 40% reduction in endogenous *Wnt4* expression at E14.5 (**Fig6I**). This confirms that mCRS4 indeed regulates *Wnt4 in vivo*, but its exact requirement and relative importance for the overall *Wnt4* expression levels is context dependent.

### A two-step model for *Wnt4/WNT4* gene activation downstream of progesterone

Although deletion of hCRS4 specifically abrogates PGR-dependent induction in human MCF7 breast cancer cells, neither mCRS4 nor hCRS4 show strong evidence of being directly bound by PGR themselves. To further investigate the connection between PGR, GRHL and *WNT4*, we performed bulk RNA-seq in the PR-responsive breast cancer cell line, T47DS^76^. Treatment with the synthetic progestin R5020 for either 4 or 24 hours revealed interesting dynamic response patterns for *PGR, WNT4* and the 7 candidate transcription factors identified earlier (**Fig7A**). Specifically, expression of *GRHL1* and *GRHL2* is rapidly induced and maintained in response to progesterone signaling. Conversely, expression of *WNT4* is initially repressed and only becomes induced after 24 hours. We validated these results in an independent experiment by qRT-PCR analysis (**Fig7B**). Expression of the negative feedback target gene *FKBP5* gradually and steadily increases expression between 4 and 24 hours of R5020 treatment. *GRHL2* expression is also subtly but consistently induced starting 4 hours following the start of R5020 treatment. In contrast, *WNT4* expression is low up until 8 hours following the start of R5020 treatment and does not increase until 24 hours after the start of PGR signaling. Combined, these data show a temporal delay in *WNT4* expression in response to progesterone, which coincides with the gradual upregulation of factors that are essential for chromatin organization of the *WNT4* locus.

**Fig 7.**
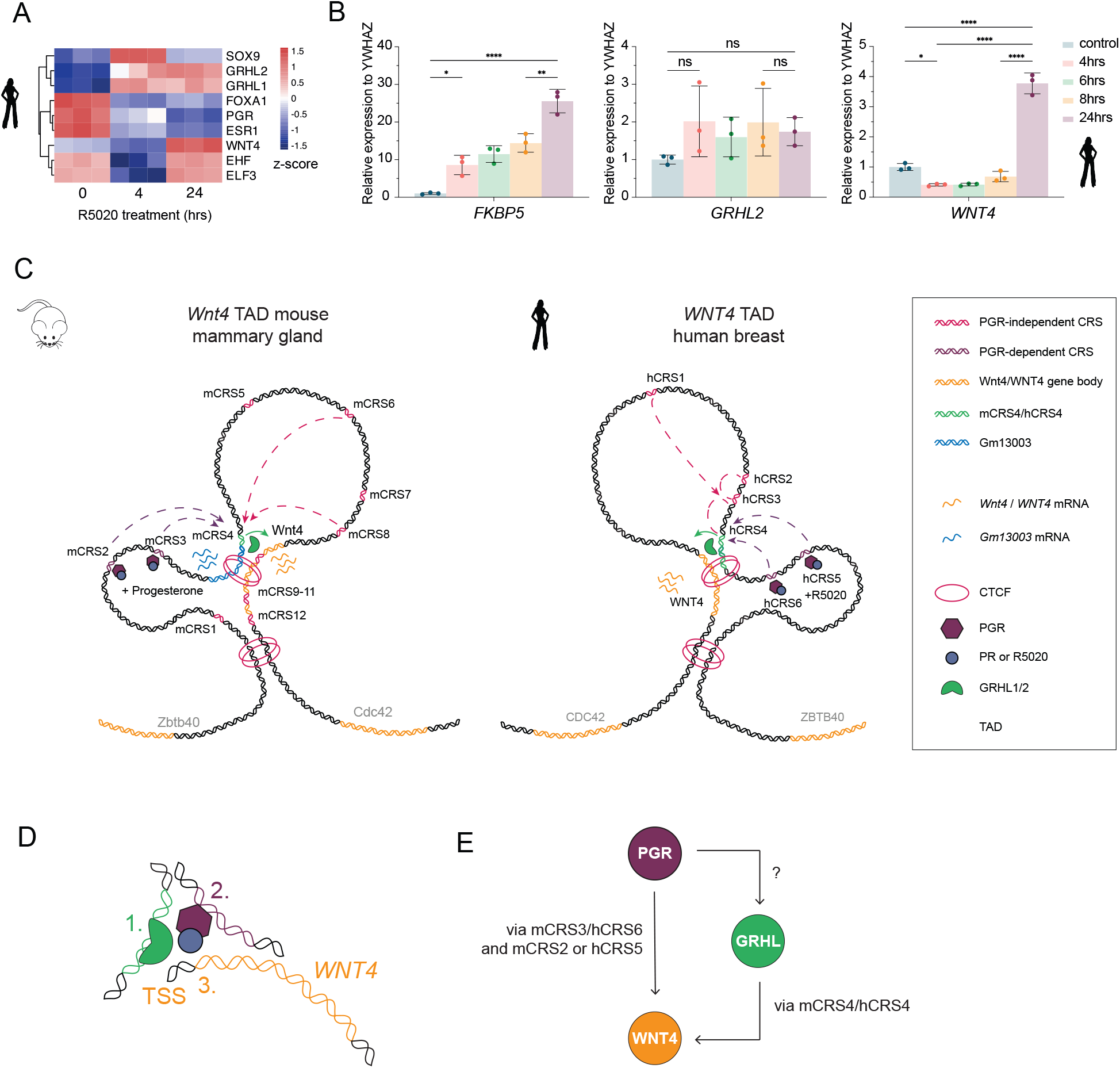
A two-step model for *Wnt4/WNT4* gene activation downstream of progesterone. A) Heatmap of bulk RNA-seq data from T47DS human breast cancer cells, showing unsupervised clustering and expression changes of 7 candidate *WNT4* regulatory transcription factors alongside *WNT4* and *PGR* in response to treatment with R5020 for either 4 or 24 hours. Expression values depicted as z-scores for n=3 replicates. B) Bar graphs of qRT-PCR data, depicting mean relative expression of *FKBP5, GHRL2* and *WNT4* after 4, 6, 8 or 24 hours of PGR signaling (R5020 treatment) in T47DS cells. Reference gene: *YWHAZ*. Datapoints: individual values for n=3 biological replicates depicted as mean fold change normalized to control. Asterisks: statistical significance (* = p < 0.05, ** = p <0.01, **** = p < 0.001, ns = not statistically significant). C) Model for 3D chromatin organization of the mouse *Wnt4* (left) and human *WNT4* (right) TAD. In the mouse mammary gland and the human breast, mCRS4/hCRS4 is a conserved CTCF and GRHL bound enhancer that loops to the *Wnt4/WNT4* promotor irrespective of *Wnt4/WNT4* expression. The mCRS4/hCRS4 enhancer subsequently functions as (part of) a distal chromatin hub on which other enhancers converge to dynamically regulate *Wnt4/WNT4* expression. D) Model for how GRHL (1, green) and progesterone-bound PGR (2, purple) converge in 3D genomic space, while binding distant regulatory elements, to jointly regulate *Wnt4/WNT4* expression (3, yellow). E) Model for a coherent feedforward loop between PGR and GRHL that explains delayed activation of *WNT4* expression in response to progesterone. PGR regulates *Wnt4/WNT4* directly (via mCRS2/mCRS3 and via hCRS5/hCRS6), while simultaneously inducing *GRHL* expression. In turn, *GRHL* enables chromatin remodeling to sustain *Wnt4/WNT4* expression via mCRS4/hCRS4.

Because gene regulation ultimately takes place in the context of 3D chromatin space, we sought to integrate our combined findings into a model for *Wnt4/WNT4* gene regulation (**Fig7C**). We propose that the overall chromatin conformation of the mouse *Wnt4* and the human *WNT4* TAD is highly similar, if not fully identical. A conserved, CTCF and GRHL bound enhancer (mCRS4 in mice, hCRS4 in humans) functions as (part of) a distal enhancer hub to coordinate PGR-dependent and independent *Wnt4/WNT4* gene expression regulation by dynamically recruiting other enhancer elements and their associated regulatory factors.

Architecturally, mCRS4/*Wnt4* and hCRS4*/WNT4* form a stable, pre-established and PGR-independent loop in the *Wnt4/WNT4* TAD. Other cis-regulatory elements, including species-specific (mCRS2/hCRS5) and conserved (mCRS3/hCRS6) PGR-responsive enhancers, are partitioned into two different clusters upstream and downstream of mCRS4/hCRS4 (**Fig7C**). Temporally, PGR-responsive gene activation is enabled by the chromatin remodeling activity of GRHL pioneer transcription factors. Ultimately, these spatial (**Fig7D**) and temporal (**Fig7E**) features converge to allow cooperative binding of PGR and GRHL2, which can come in close physical proximity through DNA folding, even though their binding sites are far removed in linear distance (**Fig7C-D**). This model also explains the weak traces (‘shadow peaks’) of PGR ChIPseq signal at mCRS4 and hCRS4 (**Fig1F, Fig3C-C’**) and supports a two-step model for *Wnt4/WNT4* gene activation downstream of progesterone.

## Discussion

In our efforts to understand tissue-specific WNT gene expression, we have uncovered a mechanistic framework for spatiotemporal regulation of WNT4 expression in the mouse mammary gland and the human breast. Our model integrates 3D chromatin accessibility and looping with the spatiotemporal binding of individual transcription factors at distinct cis-regulatory elements in the context of a complex and dynamic mammalian tissue. Specifically, we show the existence of a conserved distal regulatory chromatin hub on which multiple luminal-lineage specific enhancers and transcription factors converge to activate WNT4 in a manner that is only partially progesterone-dependent. Our working model (**Fig7C-E**) explains the joint regulation of *WNT4* by GRHL2 and PGR, including the delayed induction of *WNT4* expression in response to progesterone signaling. More broadly speaking, it redefines our view of transcriptional regulatory complexes as the combined 3D protein-DNA and protein-protein affinities.

### Structural stability and dynamic enhancer exchange

Despite the overall low sequence conservation of individual cis-regulatory elements^103,104^, the existence of a distal, regulatory chromatin hub (involving mCRS4 and hCRS4 in mice and humans, respectively) is a defining feature of the *WNT4* TAD (**Fig2A-E, Fig3A-B**). Deletion of hCRS4 in human breast cancer cells abrogates the progesterone-inducibility of *WNT4*, even though it lacks a PGR binding motif and is most likely not directly bound by PGR itself (**Fig5J**). Germline deletion of mCRS4 alone does not have a major impact on *Wnt4* expression *in vivo* (**Fig6**), suggesting rewiring or functional redundancy within the enhancer network. Such genetic compensation is not uncommon and may in fact be critical to ensure developmental robustness^96^. Taken together, we conclude that mCRS4 and hCRS4 play a key role in regulating *WNT4* gene expression in the mouse mammary gland and the human breast: they physically interact with the *WNT4* gene (**Fig2A-E, SuppFig3A**), are bound by known chromatin regulatory factors (CTCF and GRHL2, **Fig5A-E**) and form a central part of the distal upstream enhancer hub within the *WNT4* TAD to which other cis-regulatory elements are recruited to regulate *WNT4* expression (**Fig3A-B**).

It is tempting to speculate that mCRS4 also helps to stabilize the TAD boundary between *Wnt4* and *Cdc42* to prevent aberrant activation of either gene by their respective tissue-specific enhancers. *In vivo*, expression of *Cdc42* in the embryonic lung shows a minor, but statistically significant increase (10% in homozygous ΔmCRS4 embryos compared to wildtype, **Fig6J**) while expression of *Wnt4* simultaneously decreases (50% in homozygous ΔmCRS4 embryos compared to wildtype, **Fig6I**). *In vitro*, we also measure a minor effect on *Cdc42* expression in mouse mammary epithelial cell types in which mCRS4 has been knocked out (**Fig5G**).

We propose that the CTCF-bound mCRS4/hCRS4 acts as a stable docking site that can reel-in other, tissue-specific enhancers, bound by different transcription factors in different cell types as needed. As the role of CTCF in genome organization is becoming more and more defined^105–108^, this may turn out to be a general mechanism^109^. The structural basis for this could be loop extrusion facilitated via the cohesion complex, which can create the conditions for liquid-liquid phase separation to occur, by bringing distal cis-regulatory elements, the promoter and DNA-binding transcription factors in close enough proximity to interact and form a stabilized, active regulatory chromatin hub^110–113^. If loop extrusion is indeed initiated at the CTCF site in mCRS4/hCRS4, this would mechanistically explain formation of a chromatin hub centered around this conserved enhancer in the *WNT4* TAD. Indeed, components of the cohesin complex (SMC1 and RAD21) are present at the mCRS4/hCRS4 element in both human (**Fig1D’**) and mouse (**Fig2C**) cells.

Cohesin itself has no known sequence specificity. Therefore, it may be recruited to its binding sites by sequence-specific transcription factors. Interestingly, in embryonic stem cell to epiblast-like cell differentiation, GRHL2 binding motifs were found to be enriched at novel, epiblast-specific SMC1 binding sites, with GRHL2 and SMC1 showing correlated binding patterns^114^. In ovarian cancer cell lines, enrichment of RAD21 at distal enhancers was shown to be dependent on GRHL2 expression^115^. Since binding of these factors also overlaps at mCRS4/hCRS4 (**Fig1D’, Fig2C, Fig5A-D**), we propose that GRHL2 recruits the cohesin complex to this enhancer, initiating loop extrusion from this position and thereby inducing the formation of the mCRS4/hCRS4 chromatin hub. We speculate that GRHL2 first acts as a pioneer factor and subsequently as a co-activator for PGR to spatially and temporally coordinate chromatin accessibility, 3D chromatin conformation and PGR-dependent *WNT4* gene expression. This is supported by the fact that both *WNT4* and *PGR* are predominantly expressed in mature luminal breast epithelial cells, whereas *GRHL2* expression emerges earlier in luminal development (**SuppFig1C-E, SuppFig5**) and its activity is also evident in luminal progenitor cells (**SuppFig4**).

Strictly speaking, we have yet to reveal the precise functional role of mCRS4, since the knockout allele not only removes both the CTCF and the GRHL binding sites, but also the TSS of the lncRNA *Gm13003* (**Fig6A**). In mice, *Gm13003* expression correlates with *Wnt4* (**SuppFig2C-D**) and could thus contribute to *Wnt4* gene expression-regulation. As this specific lncRNA is not conserved between mice and humans, we did not follow up on its role specifically. However, we do note that *Gm13003* was reported to be upregulated >2-fold in organoids derived from mammary carcinomas compared to normal mammary organoids^116^. In the past decades, a growing body of evidence has shown that lncRNAs (and RNA in general) play an important role in gene regulation to help achieve transcriptional specificity by recruiting chromatin modifying proteins and by regulating 3D genome organization^117–122^. In many cases, the specific sequence of the cis-acting lncRNA may be less important for its regulatory function than the enhancer signature of its promoter^123–125^. We propose that *Gm13003* (or rather, its promoter as part of mCRS4) acts in a similar fashion and helps to stabilize the mCRS4 hub by binding and concentrating diffusible regulators^126–128^. This includes GRHL2, which not only directly binds to mCRS4, but which is also known to co-opt a histone methylase, KMT2C^129,130^, to prime enhancers for activation by promoting H3K4 mono-methylation to further stimulate enhancer-promoter contacts^131,132^. Other, yet to be identified sequences may play a similar role in the human *WNT4* TAD, given that here the regulatory chromatin hub also extends beyond hCRS4 (**Fig4B-D’**).

### Limitations of the study

Some technical and experimental limitations should be considered. First, while we took advantage of existing datasets and functionally validated critical findings, it is important to consider that clonal variation will exist between cell lines grown in different labs. Data obtained from primary tissues will show an even larger degree of epigenetic heterogeneity between individuals and, especially for the mammary gland, may also show dynamic changes depending on the precise developmental and reproductive stage of the tissue. Second, the *in vitro* model systems in which we can study PGR dependent processes in mammary gland cells are limited to human breast cancer cell lines such as MCF7 and T47DS. So far, we have been unable to robustly measure progesterone-dependent responses in murine cells.

Of note, our pipeline for identifying novel transcriptional regulators uses differential motif enrichment as a proxy to predict transcription factor activity. As such, we will have missed ubiquitously expressed, but nevertheless relevant, transcription factors. Additionally, because we have used ChromVAR to detect transcription factor activity, we can only identify candidates with a known and specific DNA binding motif that is included in the analysis pipeline. For example, the consensus PGR binding motif is shared with other steroid hormone receptors, which explains why it does not show up as specifically enriched in mature luminal cells in our ChromVAR analysis (**SuppFig4**). Ultimately, an unbiased experimental approach is warranted to identify locus-specific transcriptional regulatory complexes, but so far this remains challenging^133^.

### Summary and outlook: Feed forward gene regulation in the context of the 4D genome

Major efforts are still required to understand how architecture and function of the genome change in space and time (i.e. 4D). This study illustrates the continued need for dissecting tissue-specific enhancer function on a gene-by-gene basis in parallel to and complementary with genome-wide analyses aimed at determining the global organizing principles. Our working model is compatible with both PGR-dependent and - independent WNT4 expression regulation via flexible and combinatorial enhancer usage. Specifically, it offers a chromatin-level explanation (**Fig7C-D**) for the functional feedforward regulation of *WNT4* expression by PGR and GRHL (**Fig7E**). Coherent feedforward loops are present in multiple biological networks and have been shown to introduce time-sensitive delays in gene activation^134^. They have previously been proposed as a major regulatory mechanism through which nuclear hormone receptors can integrate signals^135^, with the nuclear hormone receptor (here: PGR) acting as the primary transcription factor, and one of its target genes (here: GRHL) acting as the secondary transcription factor upstream of the target gene of interest. In the case of *WNT4*, this would ensure that PGR-dependent *WNT4* gene expression is only increased when progesterone signaling is sustained, thus preventing aberrant induction due to small fluctuations in the input signal. We hypothesize that this will, indirectly, also contribute to defining a sharper WNT4 morphogen gradient in the tissue and, by extension, in shaping the WNT-responsive stem cell niche.

## Supplemental information

### Document S1: Supplemental Figures

Figures S1-S5 with supplemental legends, supplemental methods and supplemental references

### Document S2: Supplemental Files

Supp File 1: Details on conserved region 1 and 2 in mCRS4/hCRS4 (related to Fig4)

Supp File 2: Details on generation of the mCRS4 knockout mice (related to Fig6)

Supp File 3: Details on 4C *Wnt4* promoter viewpoint (related to Fig2)

### Document S3: Supplemental tables

Suppl Table 1: coordinates and conservation of mCRS and hCRS sequences

Supp Table 2: Mendelian ratios of *Wnt4*Δ*mCRS4* mice at weaning age

(related to Fig6)

### Document S4: Tables

Table 1: sgRNA sequences for mCRS4 and hCRS4

Table 2: genotyping and sequencing primer sequences

Table 3: overview of sgRNAs covering mCRS1-12

Table 4: qRT-PCR primers

Table 5: ATAC-seq primers used to generate library

Table 6: Primers used in 4C experiments

Table 7: source data for all NGS figure panels

Note:

Documents S1-S4 are uploaded to Biorxiv as supplementary files.

Document S4 is in excel format and can also be downloaded from https://osf.io/p2byu/

## Acknowledgements

We thank Selina van Leeuwen and other members of the MAD: Dutch Genomics Service & Support Provider, Swammerdam Institute for Life Sciences, University of Amsterdam for their sequencing services. We thank Rahmen Bin Ali of the Animal Model Facility (AMF) at the Netherlands Cancer Institute for technical support and our animal caretakers for taking care of the mice. We thank our colleagues of the Developmental, Stem Cell and Cancer Biology (DSCCB) group and others in the Cell and Systems Biology theme at the Swammerdam Institute for Life Sciences for discussions and feedback during the project and Amber Zeeman for technical assistance during its early stages. We thank Aalt-Jan van Dijk for critical comments on the manuscript.

We specifically thank the following BSc and MSc internship students: Roan van Scheppingen for help with setting up the 4C chromatin conformation capture protocol, Katja Klooster and Jelte Hermans for help with cloning the initial mCRS enhancer sequences and reporter constructs, Cátia Rodrigues Pereira for help with setting up the dCas9 experiments, Marius Messemaker for help with cloning and testing *Wnt4* sgRNA sequences, Muriel Wagner for help with the MCF7 luciferase assays and with generating the MCF7 CRISPR knockout cell lines, and Thomas Fleischmann for helping with characterization of the mCRS4 knockout mice.

The research was funded by the Dutch Research Council (NWO ALW VIDI 864.13.002 to RvA) and the European Union’s Horizon 2020 research and innovation program under the Marie Sklodowska-Curie grant agreement #706442 (to KEW). We also acknowledge receipt of multiple travel grants that have allowed us to present our work at international conferences (Madeleine Julie Vervoort Fonds to NH; Amsterdams Universiteitsfonds (AUF), Genootschap ter bevordering van Natuur-, Genees-en Heelkunde, and Stichting tot bevordering van het wetenschappelijk onderzoek in de biochemie to MTA).

## Author contributions

Conceptualization: RvA, KEW, YBCvdG, MTA, NH

Data curation: KEW, YBCvdG, MTA, NH, RvA

Formal analysis: YBCvdG, MTA, KEW, NH, IBH

Funding acquisition: RvA, KEW

Investigation: KEW, YBCvdG, MTA, NH, IBH

Methodology: KEW, YBCvdG, MTA, NH, RvA

Project administration: RvA, YBCvdG, KEW, NH, MTA

Resources: NKI

Software: YBCvdG, KEW, MTA

Supervision: RvA, ALvB

Validation: YBCvdG, MTA, KEW, NH, RvA

Visualization: YBCvdG, NH, MTA, KEW, RvA

Writing – original draft: NH and YBCvdG independently

wrote first drafts of the manuscript for their PhD thesis

with input from RvA, KEW and MTA.

Writing – review & editing: All authors provided input on the final manuscript. RvA and MTA prepared the manuscript for submission and publication.

## STAR Methods

### Experimental models

#### Cell culture

Mammary epithelial BC44 cells (a kind gift of Marie-Anne Deugnier, Institute Curie, Paris, France) were cultured in RPMI1640 + L-Glut (Fisher, 11973466), supplemented with 10% FBS (Gibco, cat# 10270-106) and 5μg/ml insulin (Sigma-Aldrich, cat. #I9278). HC11 mammary epithelial cells were also cultured in RPMI1640 + L-Glut with 10% FBS and 5μg/ml insulin, with the addition of 10ng/ml EGF (Peprotech, cat. #AF-100-15-B). Cells were dissociated with 0.05% trypsin and split 1:10 every 3 days. MCF7 human epithelial breast cancer cells (a kind gift from Pernette Verschure, Swammerdam Institute for Life Sciences, Amsterdam) and T47DS human epithelial breast cancer cells (a kind gift from Stieneke van den Brink, Hubrecht institute, Utrecht, The Netherlands) were cultured in DMEM + GlutaMAX (Gibco, 11584516) supplemented with 10% FBS. Cells were split 1:5−1:10 twice a week and routinely tested for mycoplasma.

#### Generation of BC44 CRISPR knockout lines

To generate a knockout of both the mCRS4 conserved *Gm13003* promoter and the conserved *Gm13003* promoter plus TSS (for coordinates of mCRS4 see **SuppTable 1**), BC44 cells were plated with a density of 100,000 cells per well in a 6-well plate. After 24hr, cells were transfected with either 1μg GFP pSGFP2-C1^159^ (Addgene plasmid #22881, a gift from Dorus Gadella, RRID:Addgene_22881, used as a transfection efficiency control), or 1μg PX459 (Addgene plasmid #62988^160^, a gift from Feng Zhang, RRID:Addgene_62988) without any sgRNAs, 1μg PX459 harbouring sgRNA1-8 (in an equimolar ratio) or 1μg PX459 harbouring sgRNA1-5 (in an equimolar ratio) using X-tremeGene^TM^ HP DNA Transfection Reagent (Sigma-Aldrich, cat. #6366236001) according to the manufacturer’s protocol (1:1 ratio). At 24hr after transfection selection was started by the addition of medium containing 1μg/ml Puromycin. At 48hr after starting the treatment, Puromycin was washed off, cells were trypsinized, counted and plated in a 96-well plate via a dilution series starting at 100cells / well to achieve single clones. Single colonies were expanded and checked for knockout of the mCRS4 or the *Gm13003* conserved promoter by genomic PCR. Genomic PCR was performed using Phusion polymerase (ThermoScientific) according to the manufacturer’s protocol. Three mCRS4 knockout clones, four promoter knockout clones and two unaffected control clones were obtained, verified by DNA Sanger sequencing and used for further experiments. See **Table 1** for sgRNA sequences and **Table 2** for primer sequences. Details on gRNA design and cloning below.

#### Generation of MCF7 CRISPR knockout lines

To knock out hCRS4 (see **SuppTable 1** for coordinates) 400,000 MCF7 cells per well were plated in a 6-well plate 24hr prior to transfection. Cells were transfected with either 2μg control plasmid (Addgene Plasmid #22881) or 1μg PX459/sgRNA1 and 1μg PX459/sgRNA2 using X-tremeGENE™ HP DNA Transfection Reagent (Sigma-Aldrich, cat. #6366236001) according to the manufacturer’s protocol (1:1 ratio). At 24hr after transfection, selection was started by adding medium containing 1μg/ml Puromycin. Selection medium was refreshed 24hr later. After another 24hr, cells were trypsinized and plated in one well of a 6-well plate without selection. After 3 days, cells were trypsinized and a single cell suspension was obtained by passing cells through a 70μm and then a 40μm filter. Single cells were plated in a 96-well plate with a density of 0.5 cells/well. Single colonies were expanded and checked for the 1500bp knockout of hCRS4 by genomic PCR using Phusion polymerase (Thermo Scientific) according to the manufacturer’s protocol. Two wildtype and two knockout clones were obtained, verified by DNA Sanger sequencing and used for further experiments. See **Table 1** for sgRNA sequences and **Table 2** for primer sequences. Details on gRNA design and cloning below.

#### Mouse strains

All animal experiments were approved by the Animal Welfare Committee of the University of Amsterdam (2013-2017: local permit) or the Centrale Commissie Dierproeven (2018-2022: AVD1110020173145, 2023-2027: AVD11100202216423). All mice were maintained on an FVB/N background under standard housing conditions in either open or filtertop cages, with 12hr light/dark cycle and *ad libitum* access to food and water. For colony maintenance, wildtype, inbred FVB/NHan®Hsd mice were purchased from Envigo.

CRISPR/Cas knockout mice were generated by the Animal Modeling Facility (AMF) of the Netherlands Cancer Institute (NKI) (permit: AVD30100202115197) on an FVB/N background using iGonad^161,162^. The genome-editing solution used to generate the knockout alleles contained Cas9 protein (IDT) and 2 gRNAs (target sequences 5’-GCTCTGGGTGACCCGTAGTTTGG-3’ and 5’-CTGCTCCTGTTATACCTAGGGGG-3’, designed using CRISPOR). Offspring was screened by PCR and Sanger sequencing. See **Table 2** for genotyping primers.

Two independent founders were used to establish the mCRS4Δ1 (1351bp deletion, removing mm10 chr4:137,163,990-137,165,340) and mCRS4Δ2 (1352bp deletion, removing mm10 chr4:137,163,991-137,165,342) lines. See **SuppFile2** for details. The two lines were maintained via heterozygous backcrosses and characterized independently. All experiments for this study were performed using F1, F2 or F3 animals. For all experiments depicted in **Fig6** and **SuppTable2**, wildtype and homozygous knockout mice were obtained from heterozygous intercrosses of mCRS4Δ1 or mCRS4Δ2 animals. Where possible, littermates were compared. Data for mCRS4Δ1 and mCRS4Δ2 were pooled, since no differences were detected between the two lines.

#### Genotyping

Standard genotyping was performed using a three-primer PCR to discriminate the wildtype (561 bp) and mCRS4 knockout (397 bp) alleles. A common forward primer (RVA3844, 5’ – GCTGCCTTCACTCTGATCAGC – 3’) was used in combination with a wildtype reverse (RVA3845, 5’ – GCAAGGGTTGGAGCAGCTACTT – 3’) and a knockout reverse (RVA3846, 5’ –CCTGGGATGGTGGTCAGAAT – 3’) primer.

Tissue samples (ear clips or tail cults) were lysed using 200μl of Direct PCR lysis reagent Viagen, 102-T) with 1% proteinase K (VWR, 0708-100MG) and incubated at 55°C overnight (O/N). Proteinase K was inactivated at 85°C for 45min, samples were spun down and 1μl of lysate was used or a standard 20μl PCR reaction with 1x Phire Green buffer (Thermo Fisher, CatNo. F527L), 2μM dNTPs (Thermo Scientific, R0181), 2μM shared forward and 1μM of each reverse primer, and 0.4μl Phire Green Hot Start II polymerase (Thermo Fisher, CatNo. F124S). PCR was performed in a Biometra TRIO cycler (30sec denaturation at 98°C, followed by 35 cycles of denaturation for 5sec at 98°C, 5sec annealing at 60°C, 10sec extension at 72°C, followed by 1min extension at 72°C). Samples were cooled to 16°C and run on a 2% agarose gel.

### Experimental Techniques

#### Cloning enhancer luciferase constructs

We cloned a minimal promoter (identical to the minimal promoter of pGL4.23 (Promega): TAGAGGGTATATAATGGAAGCTCGACTTCCAG) into the HindIII site of pGL4.20, which contains a firefly luciferase gene (*luc2*) and a puromycin resistance gene. This pGL4.20 plasmid with the minimal promoter is referred to as pGL4-minP.

Sequences for mCRS1-12 and hCRS1-6 (corresponding to the coordinates shown in **SuppTable 1**) with compatible ends were ordered as double stranded gBlocks from IDT. The default combination was XhoI (5’)/BglII (3’), but when the CRS contained one of those restriction sites, SalI (5’) and/or BamHI (3’) were chosen. We adjusted the lengths of all mouse and human enhancer sequences by adding max. 200bp to both the 5’ and 3’ ends of the core sequence that was originally called as a peak (mouse: ATAC-seq and/or ChIP-seq) or element (human: Cicero analysis).

Tubes from IDT were spun down and DNA was diluted with 0.2μm filtered water to a concentration of 10ng/μl, after which the tubes were incubated at 50°C for 20min. The pGL4-minP vector was digested with XhoI and BglII and 1X Tango buffer (Thermo Scientific). gBlocks were digested with the appropriate compatible enzymes. After digestion, gBlocks were purified using the GeneJET PCR Purification kit (Thermo Scientific, cat. #K0701). pGL4-minP vector and CRS inserts were ligated using T4 ligase (Thermo Fisher Scientific, cat. #EL0016) and T4 DNA ligase buffer in a ratio of 1:3 (vector:insert) O/N at 16°C.

#### Design and cloning of sgRNA constructs

For mouse gRNA design, CRISPOR (http://crispor.tefor.net) was used. For each enhancer, one sgRNA was designed for every 150bp. For human gRNA design, the gRNA design tool in Benchling (https://www.benchling.com) was used. All gRNA sequences are provided in **Table 1**. Cloning was done according to the Zhang lab protocol^160^. To generate knockout cell lines, gRNAs were cloned into pSpCas9(BB)-2A-Puro (PX459, Addgene plasmid #62988^160^, a gift from Feng Zhang, RRID:Addgene_62988) at the BbsI site. For the dCas9 assays, gRNAs were cloned into pSpgRNA (Addgene plasmid #47108^163^, a gift from Charles Gersbach, RRID:Addgene_47108) at the BbsI site.

#### Cloning GRHL overexpression constructs

For GRHL overexpression experiments in serum-starved BC44 cells we used our previously generated pGlomyc3.1-Grhl3 construct^164^ (Addgene #172869, RRID:Addgene_172869). Sequences of Grhl1 and Grhl2 without introns were ordered as double stranded gBlocks from IDT and cloned into pGlomyc3.1 to generate pGlomyc3.1-Grhl1 and pGlomyc3.2-Grhl2. These constructs are currently being deposited with Addgene under accession numbers #236049 and #236050.

#### Transfections for luciferase assays

BC44, HC11 (both 20,000 cells per well) were plated in a 24-well plate 24hr prior to transfection. X-tremeGENE™ HP DNA Transfection Reagent (Sigma-Aldrich, cat. #6366236001) was used for transfection according to manufacturer’s protocol (1:1 ratio). Transfections were done in triplo, using 300ng pGL4_minP (empty vector control or containing the CRS sequence), 100ng CMV-Renilla (transfection control), and 100ng eGFP (to check transfection efficiency) in each well. Cells were harvested 48hr after transfection, at which time plates were stored at -80°C until further analysis (i.e. cell lysis and luciferase assays).

MCF7 cells (100,000 cells per well) were plated in a 24-well plate 24hr prior to transfection. At 24hr after plating, culture medium was switched to DMEM + GlutaMAX (Gibco, 11584516) supplemented with 5% Charcoal Stripped Fetal Bovine Serum (Thermo fisher, A3382101). X-tremeGENE™ HP DNA Transfection Reagent (Sigma-Aldrich, cat. #6366236001) was used for transfection according to manufacturer’s protocol (1:1 ratio). Transfections were done in duplicate using 200ng pGL4_minP (empty vector or containing the CRS sequence), 100ng CMV-Renilla (transfection control), and 200ng PGRb vector (Addgene Plasmid #89130^165^, a gift from Elizabeth Wilson, RRID:Addgene_89130) in each well. For hormone treatment, MCF7 cells were refreshed 48hr after plating with medium supplemented with 5% Charcoal Stripped Fetal Bovine Serum containing either EtOH, 20nM R5020 (Promegestone) (Perkin Elmer, NLP004005MG), or 20nM R5020 and 100nM RU486 (Mifestrone) (Sigma Aldrich, 475838). Cells were harvested 72hr after transfection (24hr after treatment), at which time plates were stored at -80°C until further analysis (i.e. cell lysis and luciferase assays).

#### Transfections for dCas9 assays

BC44 and HC11 cells were plated with a density of 100,000 cells per well in a 6-well plate. After 24hr, cells were transfected with prepared pools of sgRNA (500ng total per well, divided over 4-8 gRNAs per mCRS, see **Table 3**) and dCas9-VPR (1500ng, Addgene #63798^84^, a gift from George Church, RRID:Addgene_63798) using X-tremeGENE™ HP DNA Transfection Reagent in a 1:1 ratio. Cells were harvested after 48hr, at which time plates were stored at -80°C until further analysis (i.e. RNA isolation and qRT-PCR).

#### Transfections for overexpression GRHL proteins

For overexpression of GRHL1, GRHL2 or GRHL3, 300,000 BC44 cells were plated in a 6-well plate, 24hr prior to transfection. Cells were transfected using 2μg pGlomyc empty vector (control) or 2μg of either pGlomyc-Grhl1, pGlomyc-Grhl2, or pGlomyc-Grhl3 using X-tremeGENE™ HP DNA Transfection Reagent (Sigma-Aldrich, cat. #6366236001) according to manufacturer’s protocol (1:1 ratio). At the time of transfection, the medium was switched to starvation medium containing 1% FBS. At 24hr after transfection, new starvation medium was added. At 48hr after transfection, samples were harvested for RNA isolation and qRT-PCR.

#### Transfection for siRNA knockdown of GRHL proteins

For siRNA knockdown of GRHL1/2/3, 150,000 MCF7 cells per well were plated in a 12-well plate, 24hr prior to transfection. SiRNA transfection for combined knockdown of GRHL1/2/3 was performed by mixing ON-TARGETplus siRNA SMARTpools (Dharmacon) of GRHL1/2/3 to a final concentration of 25nM for each pool in Opti-MEM and transfecting these pools using the Dharmafect transfection reagent (Dharmacon) according to the manufacturer’s protocol. At 48hr after transfection, samples were harvested for RNA isolation and qRT-PCR.

#### Dual luciferase assays

Cells were lysed in 1X Passive Lysis Buffer (50μl per well, Promega, cat# E1941)) according to manufacturer’s instructions. For the reactions, non-commercial firefly and Renilla luciferase reagents (LAR) were used^166^. Firefly LAR is composed of 200mM Tris HCl (pH 8.0), 15mM MgSO_4_, 0.1mM EDTA (pH 8.0), 25mM DTT (Fisher cat. 10792782), 1mM ATP pH 7.0 (Sigma cat. A2383), 0.2mM Coenzyme A (Sigma cat. C3144), 200μM D-Luciferin pH 6.0-7.0 (Biosynth cat. L-8200) with a final pH of 8.0. The buffer of Renilla LAR contains 25mM Na_4_P_2_O_7_ (Carl Roth cat. T883.1), 10mM NaAc, 15mM EDTA, 500mM Na_2_SO_4_, 500mM NaCl with a final pH of 5.0. The following components were freshly added to the Renilla LAR buffer before every assay: 50μM phenylbenzothiazole (Santa Cruz Biotechnology cat. sc-391075) and 4μM benzyl coelenterazine (Nanolight cat. 301-500). Firefly and Renilla luciferase activity was measured in a GloMax Navigator (Promega, cat# GM2000). The following protocol was used in the GloMax: Injection of 50μl non commercial firefly LAR, 2sec pre-measurement delay, 10sec measurement firefly reporter, injection of 50μl non-commercial Renilla LAR, 2sec pre-measurement delay, 10sec measurement Renilla reporter. Each individual Firefly luciferase values was normalized to its own corresponding Renilla luciferase value.

#### RNA isolation and qRT-PCR analysis

For all cell culture experiments, whole mammary glands, embryonic lungs, and FACS sorted primary mammary cells, RNA was isolated using Trizol (Invitrogen, cat. #15596018) according to the manufacturer’s protocol. cDNA synthesis was performed with 200ng - 4μg of RNA using SuperScript IV Reverse Transcriptase (Invitrogen, cat. #18090200) and Random Hexamers (Invitrogen, cat. #N8080127) according to manufacturer’s guidelines with the addition of RiboLock RNase Inhibitor (Thermo Scientific, cat. #EO0328). After cDNA synthesis was completed, samples were diluted 10x. qRT-PCR reactions were performed with the primers in **Table 4** using a QuantStudio 3 Real-Time PCR System. For the reactions, 5μl of diluted cDNA was added to a mix of 4μl 5X HOT FIREPol EvaGreen qPCR Mix Plus (ROX) (Solis Biodyne, cat. #08-24-00008), 2μl primers (from a 10μM stock) and 10μl nuclease-free water. For all qRT-PCR experiments, reactions were performed in technical triplicates in a 96x0.2 ml plate (BIOplastics, cat. #AB17500). The following stages were included in the reaction: 2min at 50.0°C and 15min at 95.0°C, then 40 cycles of 15sec at 95.0°C and 1min at 60.0°C, followed by the melting curve stage. Relative expression for the BC44 knockout cell lines was calculated based on 3 independent reference genes^46^ (*Ctbp1, Prdx1* and *Rpl13a*) according to Riedel et al.^158^. In all other cases, a single reference gene was used as indicated in the figure legends. For further gene expression quantification, relative gene expression was calculated according to the ΔΔCT method.

#### Carmine staining and quantification

For Carmine alum staining of mouse mammary glands, freshly isolated mammary glands were transferred to an electrostatically adherent microscopy slide. Glands were flattened by adding another microscopy slide (Carl Roth 2105.1) on top and the two glasses were stored in a 50ml tube containing Carnoy’s solution (75% EtOH (VWR 83804.360), 25% glacial acidic acid (Sigma 137130)) overnight. The next day, glands were taken out of the Carnoy’s solution, removed from the slides and added to a plastic container filled with carmine alum solution (200mg Carmine dye (Sigma C1022) and 500mg aluminum potassium (Sigma 237086) in 100ml dH2O). Glands were left shaking in Carmine solution for 24-72hr. Destaining of the glands was then performed shaking in destaining solution (70% EtOH (VWR 83804.360) containing 2ml/l 12N HCl (Sigma H1758)). Destaining solution was refreshed every 30min until the solution remained clear, samples were stored in destaining solution 24-72hr after the clear solution was reached. Next, dehydration was performed by submerging the glands for 5min each in 70, 80, 95 and 100% EtOH. Dehydrated glands were stored in Histoclear II (Brunschwig Chemie, HS-202) overnight. For imaging, glands were transferred to a microscopy slide and imaged on an Epson flatbed scanner at a resolution of 4800dpi. After initial imaging, glands were mounted on an electrostatically adherent microscopy slide using Omnimount (National Diagnostics, HS-110). For quantification, images were loaded into Fiji2 (Version 2.14.0). Fat pad outlines were manually selected using the segmented line tool. The total fat pad area was then quantified (analyze → measure). The area of the fat pad containing ductal epithelium was similarly outlined using manual tracing. The percentage fat pad filling was then calculated by calculating ductal fat pad area/ total fat pad area*100.

### NGS dataset generation

#### Bulk ATAC-seq

The 3^rd^ and 4^th^ mammary gland fat pads from 6 puberty (p35) FVB/N mice were dissected, chopped to small pieces and incubated for 2hr at 37°C in an orbital shaker in a digestion mix composed of DMEM/F12, 5% FBS, 1% Penicillin / Streptomycin, 25mM HEPES and 300U/ml Collagenase IV. Red blood cells were lysed with ACK Lysing Buffer and single cell suspensions were generated by consecutive digestion steps with Trypsin-EDTA and DNAseI. Cells were stained in HBSS containing 10% FBS with the following antibodies (eBioscience): anti-Mouse CD45-Biotin (clone 30-F11), anti-Mouse CD31-Biotin (clone 390), anti-Mouse TER-119-Biotin (clone Ter-119), anti-Mouse CD326-PE (clone G8.8), anti-Mouse CD49f-FITC (clone GoH3) and Streptavidin-APC. DAPI was used for live/dead cell discrimination. 55,000 luminal, basal and stromal cells were sorted into ice-cold 10mM HEPES containing 10% FBS using a BD FACSAria III equipped with a 100μm nozzle at 20psi. FITC: excitation with 488nm laser, emission detected using a 530/30nm bandpass filter; PE: excitation with 561nm laser, emission detected using a 582/15nm bandpass filter; DAPI: excitation with 407nm laser, emission detected using a 450/50nm bandpass filter; APC: excitation with a 633nm laser, emission detected using a 660/20nm bandpass filter. Cells were sorted with a plate voltage of 2500V using the 4-Way Purity precision mode.

ATAC-seq samples were prepared according to the protocol from Buenrostro et al.^167,168^. Briefly, freshly sorted cells were washed once in 50μl ice-cold PBS and gently resuspended in 50μl cold lysis buffer (10mM Tris-HCl, pH 7.4; 10mM NaCl; 3mM MgCl2 and 0.1% NP-40) to isolate nuclei. The transposase reaction (Illumina Nextera DNA Library Preparation Kit, FC-121-1030) was performed for 30min at 30°C and DNA was purified using MinElute PCR purification columns (Qiagen, #28004). DNA fragments were amplified using NEBNext Ultra II Q5 Master Mix (#M05445) and custom primers (see **Table 5**).

The following PCR conditions were used: 72°C for 5min; 98°C for 30sec, 8 cycles of 98°C for 10sec, 63°C for 30sec and 72°C for 1min. Libraries were purified with the MinElute PCR purification kit. An additional, single left-sided purification with AMPure XP beads (Beckman Coulter #A63880) was performed to remove primer dimers. Libraries were quantified with qRT-PCR before paired-end sequencing on an Illumina NextSeq 550 system (2 x 75bp, performed by MAD: Dutch Genomics Service & Support Provider, Swammerdam Institute for Life Sciences, University of Amsterdam). Sequencing data were processed using the ATAC-Seq pipeline of the Kundaje lab

(https://github.com/kundajelab/atac_dnase_pipelines) with default parameters and mouse genome version mm9. Processed files were visualized as BigWig in the IGV browser^144^. The data have been deposited with NCBI GEO under accession number GSE291413.

#### Bulk RNAseq of BC44 and HC11

For RNA isolation from BC44 and HC11 cells a combination of TRIzol reagent (Fisher Scientific, #15608948) and RNeasy mini kit (Qiagen, #74104), including on-column DNAse digestion (Qiagen, RNase-free DNase set #79254) was used. Briefly, cells were lysed by directly adding 1ml TRIzol to 10cm culture dishes. The homogenate was transferred to Phasemaker tubes and mixed with chloroform. After separation, the aqueous RNA layer was mixed with 1 volume of 70% ethanol and transferred to RNeasy spin columns. RNA purification and DNase digestion were performed according to manufacturer’s instructions.

All library preparation and sequencing steps were performed by MAD: Dutch Genomics Service & Support Provider, Swammerdam Institute for Life Sciences, University of Amsterdam. RNA integrity was analyzed with an Agilent RNA ScreenTape system (RIN ≥ 9.6). ERCC RNA Spike-In Mix 1 (ThermoFisher Scientific #4456740) was added to samples prior to polyA enrichment (NEBNext Poly(A) mRNA Magnetic Isolation Module (New England BioLabs) and stranded-library preparation (NEBNext Ultra II Directional RNA Library Prep Kit for Illumina and NEBNext Multiplex Oligos for Illumina (New England BioLabs). Assessment of the size distribution of the libraries with indexed adapters was performed using a 2200 TapeStation System with Agilent D1000 ScreenTapes (Agilent Technologies). The NEBNext Library Quant Kit for Illumina (New England BioLabs) was used according to manufacturer’s instructions to quantify the libraries on a QuantStudio 3 Real-Time PCR System (Thermo Fisher Scientific). Libraries were clustered and sequenced on an Illumina NextSeq 550 Sequencing System (NextSeq 500/550 Mid Output Kit (300 Cycles), 2 x 150bp).

The snakemake workflow VIPER^169^ (Visualization Pipeline for RNA-seq analysis) was used to map and align raw sequencing data to the mouse genome (mm9, STAR^170^ version 2.7.1a). Raw transcript counts were further processed using edgeR^171^ (version 3.28.0) and limma^172^ (version 3.42.1) packages with R version 3.6.2^173^. A cutoff of CPM > 0.5 in at least 2 libraries was applied to filter out genes with low counts prior to trimmed mean of M-values (TMM) normalization. The data have been deposited with NCBI GEO under accession number GSE291779.

#### Bulk RNAseq of T47DS

For RNA isolation, 700,000 T47DS cells were plated in 6-well plates containing DMEM + GlutaMAX (Gibco, 11584516) supplemented with 5% Charcoal Stripped Fetal Bovine Serum (Thermo fisher, A3382101). For 24hr induction of progesterone signaling, the medium was refreshed 24 hrs after plating and cells were treated for another 24hr with either 1nM R5020 or EtOH (as a control) added to the culture medium. For 4hr induction of progesterone signaling the medium was refreshed 44 hr after plating and cells were treated for another 4 hr with either 1nM R5020 or EtOH (as a control). All cells were then harvested and further processed simultaneously.

RNA isolation was performed using the RNeasy mini kit (Qiagen, #74104), including on-column DNAse digestion (Qiagen, RNase-free DNase set #79254) according to manufacturer’s instructions. 1μg of RNA was used as starting material for poly-A enrichment using the NEBNext Poly(A) mRNA Magnetic Isolation Module (New England BioLabs). Next, RNA-Seq libraries were generated according to the manufacturers’ protocols using the NEBNext Ultra II Directional RNA Library Prep Kit for Illumina and NEBNext Multiplex Oligos for Illumina (Unique Dual Index Primer Pairs) (New England BioLabs). Distribution of the size of the libraries with indexed adapters was assessed using a 2200 TapeStation System with Agilent D1000 ScreenTapes (Agilent Technologies). The QuantStudio 3 Real-Time PCR System (Thermo Fisher Scientific) was used for library quantification using the NEBNext Library Quant Kit for Illumina (New England BioLabs) according to manufacturer’s instructions. NextSeq 500/550 High Output Kit v2.5 (75 Cycles) (Illumina) was then used for library clustering and sequencing (75 bp) on a NextSeq 550 System (Illumina). All library preparation and sequencing steps were performed by MAD: Dutch Genomics Service & Support Provider, Swammerdam Institute for Life Sciences, University of Amsterdam.

Processing of raw sequencing files was performed on the Galaxy server^174^. Quality control on FASTQ files was performed using MultiQC^175^, followed by removal of overrepresented sequences by read trimming with Trimmomatic^176^ using a list of selected adapters. Trimmed reads were then mapped to the human reference genome (version GRCh38.p14) using HISAT2^177^. Mapping quality was inspected using SAMtools^178^ stats and visualized using MultiQC. HTSeq count^179^ was then used to create a count matrix. Further differential gene expression analysis was performed using DESeq2^180^ in R studio (version 2023.06.1). The data have been deposited with NCBI GEO under accession number GSE291778.

#### 4C circular chromatin conformation capture

A 636bp DNA fragment located 650bp upstream of the *Wnt4* TSS (details in **SuppFile3**, corresponding coordinates of the *Wnt4* transcript in mm9 chr4:136833550-136852694) was used as the viewpoint. 4C-seq was performed as previously described^80,181^, with some modifications. Briefly, 1x10^6^ BC44 or HC11 cells were crosslinked with 2% (v/v) formaldehyde in PBS/10% FBS for 10min at RT in 15cm cell culture dishes while shaking. Cells were isolated by using a cell scraper, lysed for 10min in lysis buffer and homogenized with a dounce-homogenizer. Nuclei were digested with 200U DpnII (NEB) in DpnII buffer (NEB) complemented with 0.3% SDS and 2% Triton X-100 for 4hr, an additional 200U DpnII O/N and another 200U DpnII for 4 hr the following day at 37°C while shaking. The first ligation was performed with 100U T4 DNA ligase O/N at 16°C in a total volume of 15ml. Following decrosslinking and phenol/chloroform extraction, the DNA was digested with NlaIII in Cutsmart buffer (NEB) O/N at 37°C while shaking. After digestion a second ligation was carried out with T4 DNA ligase (Thermo Scientific) in a total volume of 15ml O/N at 16°C. DNA was isolated via phenol/chloroform extraction and purified with the ChIP DNA Clean & Concentrator Kit (Zymo Research).

50-100ng DNA template per reaction and 1μg in total was used to amplify PCR-specific libraries for the Wnt4 promoter viewpoint with the Expand Long Template PCR system (Sigma) using the primers listed in **Table 6**. Successful reactions were pooled and purified with the High-pure PCR Product Purification Kit (Sigma). Equal amounts of libraries were combined, and all library preparation and sequencing steps were performed by MAD: Dutch Genomics Service & Support Provider, Swammerdam Institute for Life Sciences, University of Amsterdam. For processing raw 4C-seq data, the publicly available 4C mapping pipeline for mapping (mm9) and filtering 4C data was used (https://github.com/deWitLab/4C_mapping) using default settings. The R package PeakC^182^ was used to analyze processed triplicates of 4C-seq data and identify significantly interacting regions within a window size of 400kb (https://github.com/deWitLab/peakC). The data have been deposited with NCBI GEO under accession number GSE291467.

### NGS dataset analyses

#### Source data

A complete list of datasets that were generated and/or reanalyzed for this study, with a link to individual figure panels, is provided in **Table 7**.

#### scRNA-seq data visualization

Processed scRNA-seq data was visualized with the CellxGeneVIP python package^137^, which expands upon CellxGene^183^ and offers more flexible data visualization. scRNA-seq datasets were imported as .h5ad files. Expression of genes of interest was either embedded in a UMAP plot annotated on cell type or displayed as a Dot Plot annotated on cell type with an expression cutoff for cell fraction of 1.

#### Visualization of epigenetic data

ChiaPET datasets were either formatted as .hic file and analyzed in JuiceBox^148^, or as .BigWig file and displayed as genomic browser tracks in the IGV browser^144^. Hi-C and micro-C datasets were visualized in the 3D genome Browser^138^. ChIP-seq, ATAC-seq and DNAse-seq was formatted as either .BigWig or .bed files and displayed as genomic browser tracks in the IGV browser. The mouse mammary gland scRNA-seq differentiation trajectory from **SuppFig1D** was generated using data from Bach et al.^184^ via https://marionilab.cruk.cam.ac.uk/mammaryGland/.

#### Co-accessibility Cicero visualization

We acquired Cicero co-accessibility scores for mouse mammary gland cells from NCBI GEO (GSE125523)^85^. The Cicero R package^86^ was used to visualize Cicero Connections. We assigned the *Wnt4* and *Gm13003* promoter based on Cicero Connection data from GSE125523 and generated genome browser-style plots with the *plot_connections()* function. We restricted the connections visualized to the genomic coordinates of the *Wnt4* TAD and used a co-accessibility cutoff of 0.15 and connection width of 0.5.

#### Peak2gene & co-accessibility correlation analysis

Peak2Gene infers linkage by linking co-accessibility in individual cells (scATAC-seq) to gene expression (scRNA-seq). Peak2gene correlation analysis was performed to identify putative regulatory relationships by correlating peak accessibility to imputed gene expression across scATAC-seq metacells. This procedure was performed by ArchR’s *addPeak2GeneLinks()* function with reducedDims set to ‘IterativeLSI’ and dimsToUse set to ‘1:30’. We set the correction cutoff at 0.2 and the resolution at 10. Gene expression in scATAC-seq was imputed after labelling by scRNA-seq by multiplying the scRNA-seq expression values by the anchor weights matrix defining the association between each scATAC-seq cell and each anchor. Next, low-overlapping aggregates of scATAC-seq cells were generated via a k-nearest neighbor procedure in the latent space to reduce noise and to ensure robust correlations in the features. Aggregates showing >80% overlap with any other aggregate were removed to reduce bias. The results were used to correlate the accessibility of every peak to the imputed expression of every gene on the same chromosome using an implementation of fast feature correlations in C++ using the Rccp package implemented by the ArchR R package^150^.

To map and plot the Peak2Gene and co-accessibility loops linearly to the genome, we used the *plotBrowserTrack()* with ‘groupBy’ set to the scRNA-seq clusters to cluster the pseudobulk ATAC tracks by cell type, and used the *getCoAccessibility()* function to retrieve the co-accessibility loops. Co-accessibility was performed by ArchR’s *addCoAccessibility()* function with reducedDims set to ‘IterativeLSI’ and dimsToUse set to ‘1:30’. We set the correction cutoff at 0.2 and the resolution at 10.

Since the resolution of the identified hCRSs by peak2gene and co-accessibility correlation was set to 10bp we expanded the enhancer candidate left and right by 500bp to achieve roughly 1000bp hCRSs, to match our prior size selection for mCRS1-12. Peak2gene analysis was performed with the integrated pseudo-scRNA-seq profile from Bhat-Nakshatri et al.^90^.

#### Single cell ATAC quality control

The protocol for single cell ATAC processing and further downstream analysis using ArchR was adapted from the tutorial at archr.com^150^ and from Kumegawa al.^185^. We obtained a list of unique ATAC-seq fragments with associated barcodes for 2 human breast samples from NCBI GEO (GSE184462)^87^. The list of unique fragments per barcode was read into the R package ArchR^150^ to perform quality control and doublet removal for each set individually. To enrich for cellular barcodes, we took advantage of the bimodal distributions in log10(TSS enrichment +1) and in log10(number of unique nuclear fragments) characterizing two different population of barcodes (cellular and non-cellular).

We manually set the barcode cutoffs thresholds for log10(TSS enrichment+1) at 4 and log10(number of unique nuclear fragments) at 1000. Only barcodes above these estimated thresholds in both metrics were kept as cellular barcodes for doublet detection. Doublet enrichment scores were calculated for cellular barcodes using ArchR’s *addDoubleScores()* with the knnMethod set to “UMAP”. Cellular barcodes with doublet enrichment scores >1 were marked as potential doublets and subsequently removed based on the filterRatio parameter of ArchR’s *filterDoublets()* function. We manually checked *WNT4* chromatin accessibility in both samples. At this stage we could not detect *WNT4* gene accessibility in patient sample JF1NV (GSM5589381). Therefore, this sample was dropped from further downstream processing and we continued our downstream analysis with patient sample IOBHL (GSM5589380).

#### Single cell ATAC-seq quantification

We used the R package ArchR^150^ to construct an initial feature matrix of 500bp genomic tiles across all cells in patient sample IOBHL. To reduce dimensions of the genomic tile features, we adopted the iterative latent semantic indexing^186–188^ (LSI) procedure implemented in the ArchR R package by using the *addIterativeLSI()* function. Briefly, this procedure performs term frequency-inverse document frequency (TF-IDF) normalization to upweight more informative features followed by an initial LSI reduction on the top accessible tiles. Graph-based Louvain clustering is used to identify low resolution clusters in which feature counts are summed across all cells in each cluster to identify the top 25,000 most variable features across clusters. This procedure was iterated once more by imputing the top 25,000 most variable tiles from iteration 1 as the top accessible tiles in iteration 2. The iterative LSI procedure computed 30 LSI dimensions that were then further collapsed into two dimensional UMAP embedding using ArchR’s *addUMAP()*. We used MAGIC^189^ to impute gene scores by smoothing signal across cells via the *addImputeWeights()* function. scATAC UMAP plots were generated in ArchR using the *plotEmbedding()* function.

#### Integration of scATAC-seq and scRNA-seq

Before transferring labels from scRNA-seq to scATAC-seq, gene activity scores were inferred in scATAC using ArchR’s *addGeneScoreMatrix()* function. Briefly, this method uses the following features to estimate gene activity: 1) fragment counts mapping to the gene body, 2) an exponential weighting function to giver higher weights to fragments counts closer to the gene and lower weights to fragment counts farther away from the gene, and 3) gene boundaries to prevent the contribution of fragments from other genes. To integrate scATAC-seq with scRNA-seq, we used the *addGeneIntegrationMatrix()* function with a RangedSummarizedExperiment object as input. This integration works by directly aligning cells from scATAC-seq with cells from scRNA-seq by comparing the scATAC-seq gene score matrix with the scRNA-seq gene score matrix via the *FindTransferAnchors()* function from the Seurat package^190^.

To assign cluster identity we integrated human breast scRNA-seq from Tabula Sapiens^88^. To further analyze pseudo-scRNA-seq profiles, human breast data from both Tabula sapiens and Bhat-Nakshatri et al.^90^ were added separately for each scATAC cell. The ArchR R package requires a singleCellExperiment (sce) object as input for scRNA-seq integration. To convert annData (.H5ad) to a sce file we used *readH5AD()* function from the Zellkonverter R package^191^. To convert Seurat objects to sce files we used the *as*.*SingleCellExperiment()* function from Seurat^190^.

Once scATAC-seq cells had received a cell type subcluster label, pseudo-bulk replicates were generated for each inferred cell type subcluster and pseudo-bulk peak calling was performed within each inferred cell type subcluster using MACS2^192,193^. ArchR’s default iterative overlap procedure was used to merge all peak calls into a single peak by barcode matrix across all cellular barcodes in each sample. Pseudo-bulk ATAC-seq coverage patterns with cell type clusters were converted to .BigWig by using ArchR’s *getGroupBW()* function and genomic browser tracks were displayed using the IGV browser^144^.

#### ChromVAR deviations enrichment

ChromVAR^89^ is an R package designed to predict enrichment of transcription factor binding motifs (a proxy for activity) on a per-cell basis from sparse chromatin accessibility data. Briefly, ChromVAR calculates motif enrichment for each individual cell (belonging to a specific cluster or – as in this study – cell lineage) compared to the total population of cells in the dataset (i.e. all cell clusters or cell lineages). ChromVAR is integrated in ArchR and provides two primary outputs: 1) ‘deviations’ (a bias-corrected measurement of how far the per-cell accessibility of a given feature (i.e. motif) deviates from the expected accessibility based on the average of all cells), and 2) ‘z-score’ (the deviation score for each bias-corrected deviation across all cells). The absolute value of the deviation score is correlated with the per-cell read depth.

ArchR subsamples sub-matrices of 5,000-10,000 cells independently to enable scalable analysis with chromVAR. First, we added background peaks by the *addBgdPeak()* function. This function selects background peaks based on chromVAR function *getBackgroundPeak()* which samples peaks based on similarity in GC-content and number of fragments across all samples using the mahalanobis distance. Next, we computed the per-cell deviations across all motif annotations with the *addDeviationsMatrix()* function. Transcription factor activity z-scores were superimposed on scATAC UMAP clusters by the *plotEmbedding()* function.

#### Identification of 44 candidate regulatory transcription factors

We used ArchR to correlate the ChromVAR transcription factor deviation scores to the gene expression of the transcription factor itself. Gene expression was obtained from human breast scRNA-seq datasets from both Tabula Sapiens^88^ and Bhat-Nakshari et al.^90^. Deviant transcription factor motifs were obtained by ChromVAR. This data was averaged by cell cluster via the *getGroupSE()* function and subsetted to obtain the deviation z-scores. We identified the maximum delta in z-score between all clusters to help stratify motifs based on the degree of variation observed across clusters. To identify transcription factors whose motif accessibility correlates with their own activity, we used the *correlateMatrices()* function. As input we used the ‘MotifMatrix’ (contain ChromVAR deviation z-scores) and ‘GeneIntegrationMatrix’ (containing gene expression information). These correlations were determined across many low-overlapping cell aggregates in the lower dimension Iterative LSI space. Transcription factors whose correlation between motif and gene expression is > 0.5, with an adjusted p-value < 0.01 and a maximum inter-cluster difference (maxDelta) in deviation z-score in the top quartile were considered positive regulators. The 44 candidate transcription factor regulators were plotted with *ggplot()*.

#### Transcription factor ID pipeline to shortlist 7 candidate regulators

We cross-referenced our list of 44 candidate positive transcription factor regulators with human breast scRNA-seq from Tabula Sapiens^88^ to determine in which cell types these positive regulators are enriched. Gene expression was visualized by dot plot in the CellxGene VIP python package^137^. Positive regulators that were enriched in the luminal lineage (LP and ML cells) but not in basal cells (or other cell types) were selected. Gene accessibility (by gene activity score from scATAC-seq), gene expression (by scRNA-seq) and transcription factor activity (by ChromVAR) was determined in the ArchR R package and superimposed on scATAC UMAP clusters by the *plotEmbedding()* function. To find experimental evidence of transcription factor binding by ChIP-seq (data shown in **SuppFig4**) we compiled peak called regions (by MACS2^192,193^) from publicly available datasets from ChIP-Atlas^194,195^ (included in **Table 7**). For positive transcription factor candiate regulators for which breast tissue data was available (FOXA1, PGR, ESR1 and GRHL1 and 2), we integrated peak called regions with a MACS2 cutoff of 100 to determine peak enrichment in scATAC clusters by deviation z-score in a similar manner as motif enrichment. Bed files containing peak called regions were loaded into ArchR by the *addArchRAnnotations()* function. The deviation z-score for ChIP peak-called regions was determined with the *addDeviationsMatrix()* function. ChIP-seq binding z-scores were superimposed on scATAC UMAP clusters by the *plotEmbedding()* function.

#### Statistics

Statistical testing was included as part of the bioinformatics workflows in R and Python as described for each individual method. For all qRT-PCR assays, statistical testing was performed in GraphPad prism (10.0.0). For comparisons of two or more conditions over multiple timepoints, a 2-way ANOVA followed by a Turkey’s multiple comparison test was performed to calculate p-values. For comparisons of three or more conditions an one-way ANOVA followed by a Turkey’s multiple comparison or a Šídák’s multiple comparison test were performed to calculate p-values. For experiments involving only two comparisons, an unpaired student’s *t*-test was conducted to calculate p-values. To determine Mendelian inheritance (**SuppTable 2**) of the mCRS4 knockout allele, a Chi-square test was performed in Excel. In all figures, asterisks indicate statistical significance: * = p < 0.05, ** = p <0.01, **** = p < 0.001, ns = not statistically significant.

#### Image elements

The female silhouette is from https://www.pngegg.com/en/png-ppaqj and is free for non-commercial use. The mouse icon was downloaded from the free icons library (https://icon-library.com/icon/mouse-animal-icon-3.html.html> Mouse Animal Icon # 52499). Both were traced and modified in Adobe Illustrator.

#### Software and online resources

| Resource | Link | Use |
| --- | --- | --- |
| ArchR<sup>150</sup> | https://www.archrproject.com/ | Multi-omics analysis |
| Cicero<sup>86</sup> | https://cole-trapnell-lab.github.io/cicero-release/ | ATAC-seq analysis |
| 4C mapping | https://github.com/deWitLab/4C_mapping | 4C analysis |
| PeakC<sup>147</sup> | https://github.com/deWitLab/peakC | 4C analysis |
| ZellKonverter | https://github.com/theislab/zellkonverter | scRNA-seq analysis |
| Seurat | https://satijalab.org/seurat/ | scRNA-seq analysis |
| ATAC-seq pipeline | https://github.com/kundajelab/atac_dnase_pipelines | ATAC-seq analysis |
| MACS2/3 | https://github.com/macs3-project/MACS | ChIP-seq analysis |
| BWA | https://github.com/bwa-mem2/bwa-mem2 | Data conversion |
| Bedtools | https://bedtools.readthedocs.io/en/latest/ | Data conversion |
| UCSC conversion tools | https://hgdownload.soe.ucsc.edu/admin/exe/ | Data conversion |
| TRANSFAC | https://genexplain.com/transfac/ | Motif analysis |
| Nucleotide BLAST | https://blast.ncbi.nlm.nih.gov/ | Sequence conservation |
| 3D Genome Browser | http://3dgenome.fsm.northwestern.edu/ | Hi-C analysis & visualization |
| ChIP Atlas | https://chip-atlas.org/ | Data source, see Table 7 |
| ENCODE | https://www.encodeproject.org/ | Data source, see Table 7 |
| NCBI GEO | https://www.ncbi.nlm.nih.gov/geo/ | Data source, see Table 7 |
| Galaxy server<sup>174</sup> | https://usegalaxy.eu | RNAseq analysis |
| MultiQC<sup>175</sup> | https://usegalaxy.eu | FASTQC |
| Trimmomatic<sup>176</sup> | https://usegalaxy.eu | Adapter trimming |
| HISAT2<sup>177</sup> | https://usegalaxy.eu | Read mapping |
| SAMtools<sup>178</sup> | https://usegalaxy.eu | Mapping QC |
| HTSeq count<sup>196</sup> | https://usegalaxy.eu | Generate count matrix |
| DESeq2<sup>180</sup> | R package | RNAseq analysis |
| Anaconda | https://www.anaconda.com/ | Python analysis |
| Rstudio | https://posit.co/ | R analysis |
| CellxGeneVIP<sup>137</sup> | https://github.com/interactivereport/cellxgene_VIP | scRNA-seq visualization |
| JuiceBox<sup>148</sup> | http://www.aidenlab.org/juicebox/ | ChiaPET visualization |
| Lineage trajectory<sup>184</sup> | https://marionilab.cruk.cam.ac.uk/mammaryGland/ | scRNA-seq visualization |
| IGV browser | https://software.broadinstitute.org/software/igv/ | Multi-omics visualization |
| GraphPad Prism | https://www.graphpad.com | Data visualization<br>Statistical analysis |
| Fiji2 (Version 2.14.0)<sup>197</sup> | https://imagej.net/software/fiji/ | Fat pad filling quantification |
| Adobe Illustrator | https://www.adobe.com/nl/products/illustrator.html | Assemble figures |

## Resource availability

### Lead contact

Further information and requests for resources and reagents should be directed to and will be fulfilled by the lead contact, Renée van Amerongen.

### Materials availability

The following plasmids generated in this study are being deposited to Addgene:

pGlomyc3.1-Grhl1 (Addgene #236049, pGlomyc3.2-Grhl2 Addgene #236050).

All gRNA and enhancer constructs are available upon request.

Sperm of the mCRS4 knockout mice has been frozen and is stored at the Animal Modeling Facility (AMF) of the Netherlands Cancer Institute.

### Data and code availability

All datasets that are composed of standardized data types will be made available via a public repository prior to publication:

|  |  |  |  |
| --- | --- | --- | --- |
| ATAC-seq | NCBI GEO | GSE291413 | status: public |
| 4C-seq | NCBI GEO | GSE291467 | status: public |
| RNA-seq mouse | NCBI GEO | GSE291779 | status: public |
| RNA-seq human | NCBI GEO | GSE291778 | status: public |

No new code was written for this study.

