## Supplementary Documents S1-S3 for "GRHL and PGR control WNT4 expression in the mammary gland via 3D looping of conserved and species-specific enhancers"

### Document S1

**Supplementary Figures and legends**

**Supplementary Methods**

**Supplementary References**

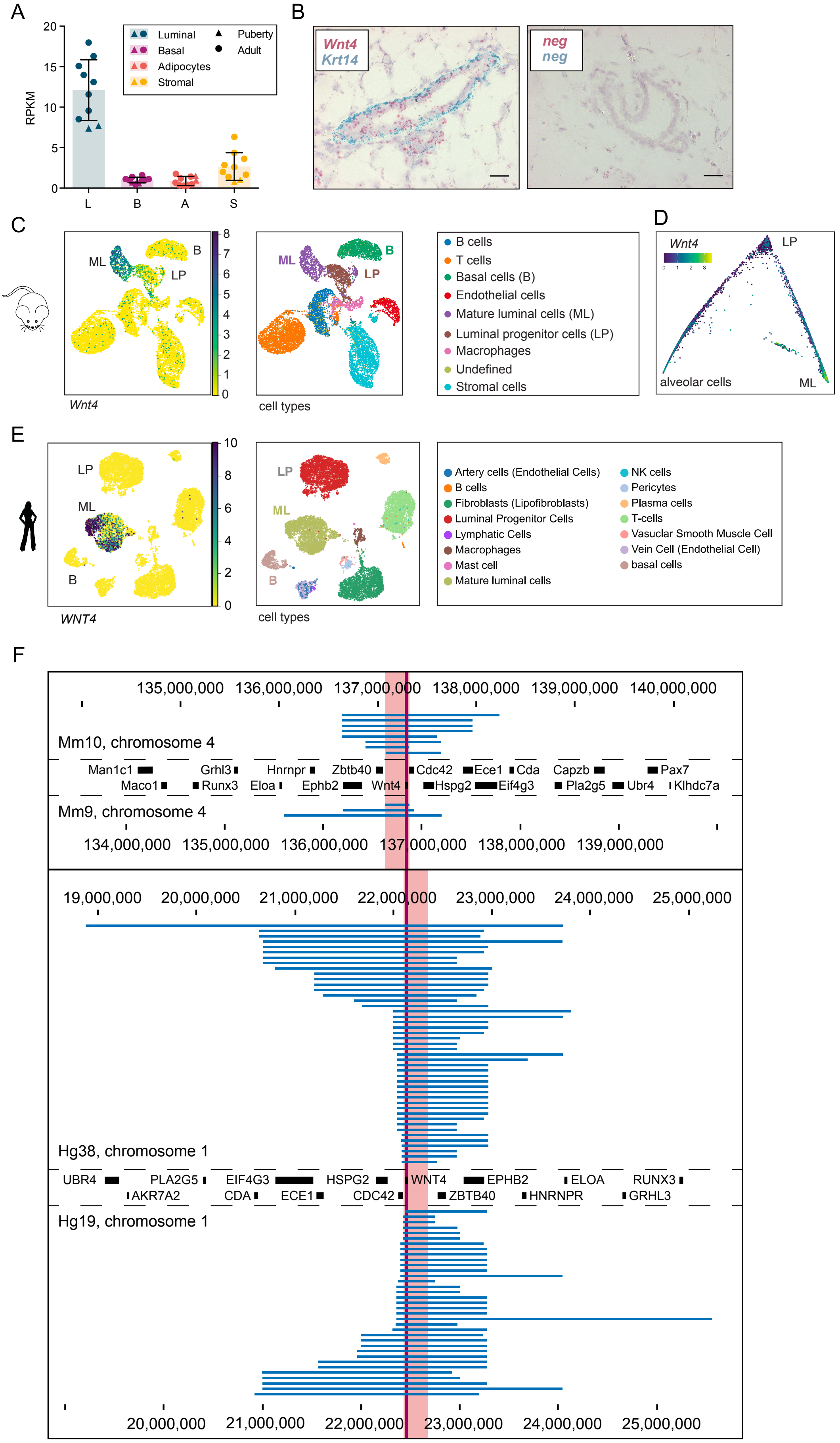

**SuppFig1. Comparison of murine *Wnt4* and human *WNT4* expression and genomic organization**
*(legend continues on next page)*

A) Bar graph showing expression of *Wnt4* measured by RNA-seq in different primary mouse mammary gland cell types isolated by FACS sorting. L = luminal, B = basal, A = adipocytes, S = stromal. RPKM = reads per kilobase per million mapped reads. Data are from FVB/N mice, published in Heijmans & Wiese et al.[1] (Fig6A in that paper depicts the same data for the n=8 adult samples split for the different estrous cycle stages, here shown pooled and supplemented with the n=2 puberty samples also described in that study).
B) RNAscope on FFPE tissue sections showing *Wnt4* (red) expression in the mammary gland of adult FVB/N mice (n=2, representative example shown). *Krt14* (blue) probes were used to mark the basal cell layer of the epithelial ducts. Nuclei were counterstained with hematoxylin. Negative controls (neg) demonstrate specificity of the staining. Scale bars are 20 μm.
C) UMAP plot displaying mouse mammary gland scRNA-seq data from Tabula Muris Senis[2], showing that *Wnt4* is predominantly expressed in mature luminal (ML) cells. Data visualization in CellxGeneVIP[3].
D) Map displaying scRNA-seq data from Bach et al.[4], showing *Wnt4* expression along the differentiation trajectory of the luminal lineage in the mouse mammary gland. LP = luminal progenitor. ML = mature luminal.
E) UMAP plot displaying human breast scRNA-seq data from Reed et al.[5], showing predominant *WNT4* expression in ML cells. Data visualized on the CellxGene website.
F) Definition of the mouse *Wnt4* and human *WNT4* TAD by aligning public Hi-C datasets, using the coordinates available in the 3D Genome Browser for mouse and human genomes[6–15] (as on 15 Feb 2019). TAD predictions (also shown in Fig1C) were called by the 3D Genome Browser[16]. *Wnt4/WNT4* gene highlighted in dark red. Coordinates of the selected TAD region (light red shading): chr4:136,625,000-136,875,000 in mice (mm9) and chr1:22,104,249-22,421,001 in humans (hg38). Only a selection of annotated genes is depicted, because of space restrictions. Note that orientation of this genomic region is inverted in human versus mouse.

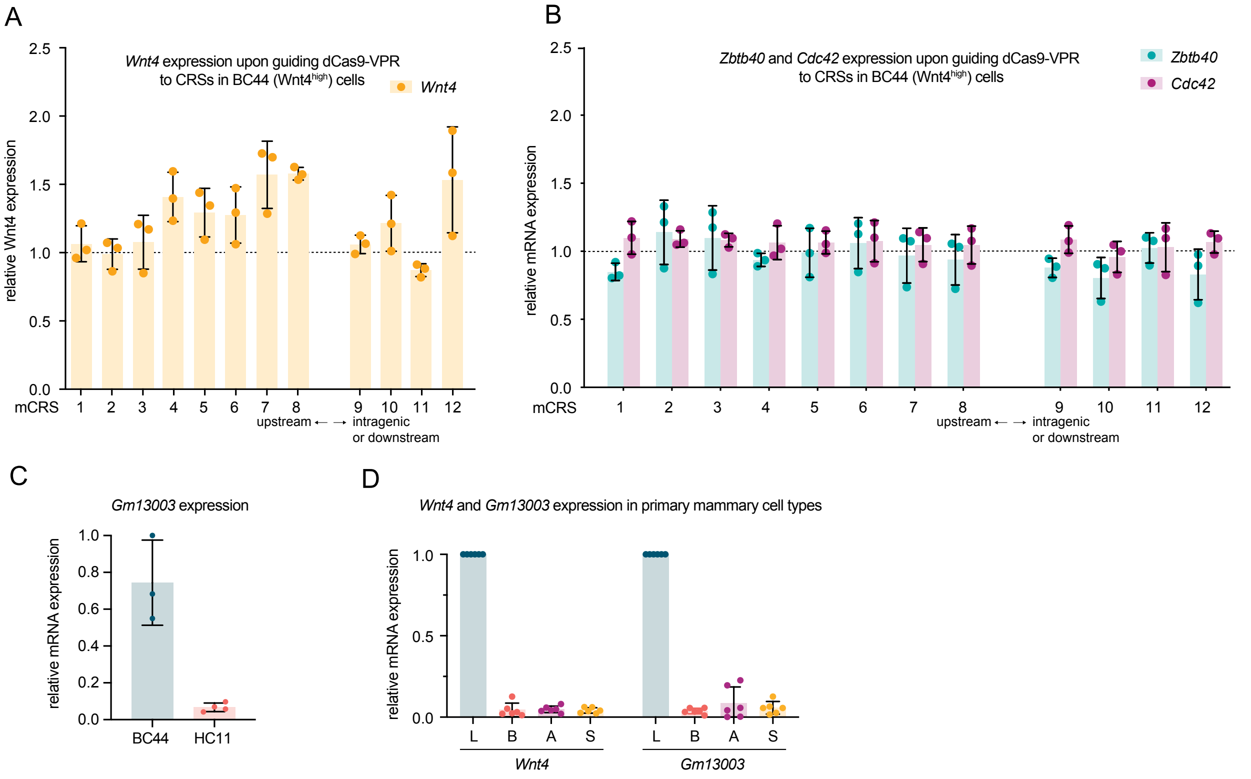

**SuppFig2. *Gm13003* expression correlates with *Wnt4* expression**
A-B) Bar graphs showing relative *Wnt4* (A, yellow), *Zbtb40* (mint green) or *Cdc42* (purple) expression measured by qRT-PCR in BC44 (Wnt4$^{high}$) cells, after guiding dCas9-VPR to individual mCRS elements. mCRSs are ordered along the x-axis according to their genomic location. The control without gRNAs was set to 1 and used to normalize all other values. *Ctbp1* was used as a reference gene[17]. Biological replicates (n=3) shown as individual data points, bars indicate the mean. Error bars: SD. This figure complements Fig2G-H, which shows the same experiment performed in HC11 (Wnt4$^{low}$) cells.
C) Bar graph portraying relative *Gm13003* expression in BC44 (Wnt4$^{high}$) and HC11 (Wnt4$^{low}$) cells as measured by qRT-PCR. *Ctbp1* was used as a reference gene[17]. The value of the highest expressing sample was set to 1 and used to normalize all other values. Data from n = 3 (BC44) or 4 (HC11) biological replicates are shown as individual data points, bars indicate the mean. Error bars: SD.
D) Bar graphs depicting relative *Wnt4* and *Gm13003* expression as measured by qRT-PCR in different mammary gland cell populations isolated from 6 adult C57/Bl6 mice. Luminal (L), Basal (B) and Stromal (S) cells were isolated using FACS and adipocytes (A) were isolated via differential centrifugation, as previously described in Heijmans & Wiese et al.[1]. *Ctbp1* was used as a housekeeping gene[17]. Values were normalized to the expression measured in luminal cells, which showed the highest expression. Datapoints represent n=6 individual mice, bars depict the mean expression. Error bars: SD.

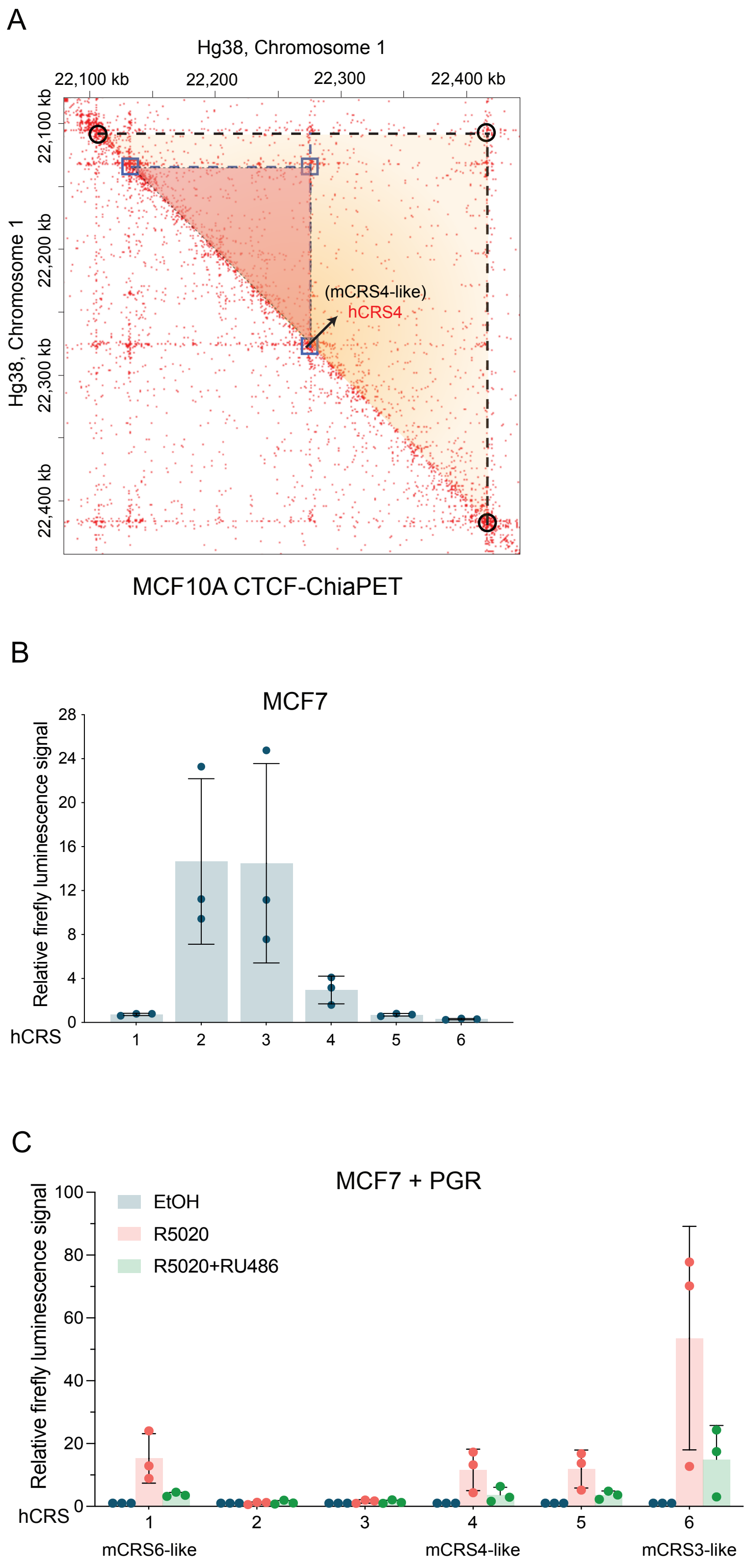

**SuppFig 3. PR-dependent and independent enhancer activity of hCRS elements**
A) CTCF-ChiaPET interaction map of the *WNT4* TAD (yellow triangle) in MCF10A cells[13,18]. Data visualized in JuiceBox[19] and aligned to hg38. Black circles indicate the *WNT4* TAD contacts. Purple squares (and red triangle) highlight the intra-TAD loop involving hCRS4.
B) Bar graph showing baseline enhancer activity of hCRS1-hCRS6 measured by transient dual luciferase reporter assays in MCF7 cells. Activity is expressed relative to an empty vector control. Datapoints show the individual values for n=3 independent biological experiments. Bars depict the mean. Error bars: SD.
C) Bar graph showing progesterone-dependent enhancer activity of hCRS1-hCRS6 measured by transient dual luciferase reporter assays in MCF7 cells transiently transfected with *PGR*. Activity in response to R5020 (red) or R5020+RU486 (green) treatment is expressed relative to the control (EtOH, blue) for each individual hCRS element. Datapoints show individual values for n=3 independent experiments. Bars depict the mean. Error bars: SD.

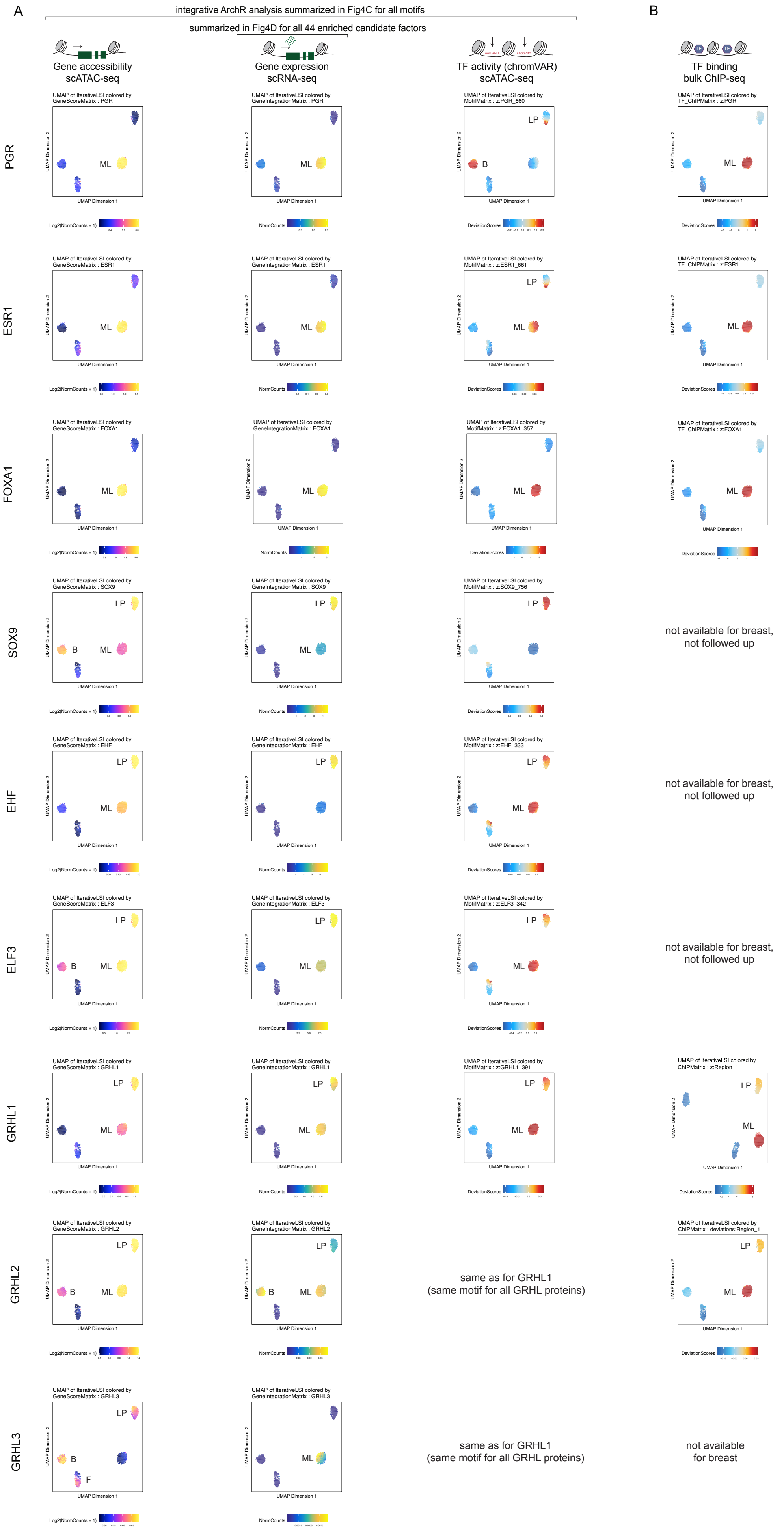

**SuppFig4. Identification of candidate transcriptional regulators of *WNT4* in the human breast**
A-B) UMAP plots depicting different cell clusters of the human breast. A) First and third column: scATAC-seq data from Zhang et al.[20], showing gene accessibility (first column) and predicted motif-based transcriptional activity (CHROMVAR[21], third column). Second column plots gene expression from scRNA-seq data by Bhat-Nakshatri et al.[22], showing gene expression enrichment in different cell clusters. B) Plots information from bulk ChIPseq data from ChIP atlas (see Table 6), showing enriched transcription factor binding in different cell clusters. Data was analysed and visualised in ArchR[23]. Cell clusters: B = basal, LP = luminal progenitor, ML = mature luminal. A selection of transcription factors is shown from top to bottom: PGR, ESR1, FOXA1, SOX9, EHF, ELF3, GRHL1, GRHL2 and GRHL3. Based on SuppFig4A and SuppFig5, we narrowed done the list of 44 candidate regulatory factors (shown in Fig4C-D) down to a final list of 7. Based on the enriched ChIP-seq binding displayed in B), we further focused on GRHL proteins.

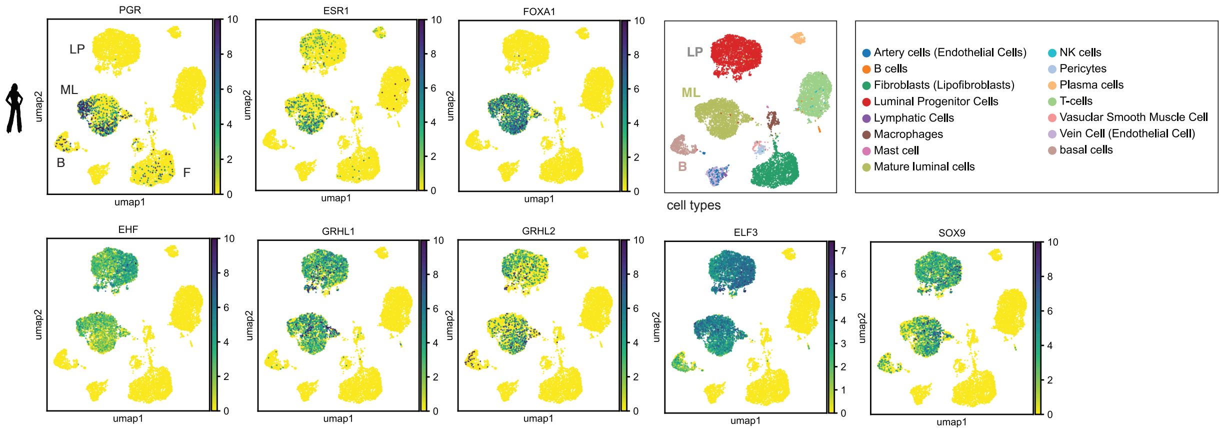

**SuppFig 5. Expression patterns of candidate transcriptional regulators of *WNT4* in the human breast**
UMAP plots displaying annotated clusters and gene expression of candidate transcription factors in human breast scRNA-seq data from Tabula Sapiens[24]. Top row: known lineage factors (PGR, ESR1 and FOXA1) and legend. Bottom row: novel candidates (EHF, GRHL1, GRHL2, ELF3, SOX9). scRNA-seq was visualised in CellxGeneVIP[3].

#### Supplementary methods

*FACS primary mammary cells*

*For SuppFig1A and SuppFig2D:* Freshly isolated mammary glands from FVB/N mice were minced and enzymatically digested for 2 hours in 9.2 ml DMEM/F12, 5% FCS, 1%P/S, 25 mM HEPES (Gibco, cat. #15630056) and 300 U/ml Collagenase IV (Gibco, cat. #17104019) (10 ml per mouse and 4 glands). Adipocytes were transferred to TRIzol LS (Invitrogen, cat. #10296028) and stored at -80°C. After resuspension in a mix (1:3) of HBSS (Gibco) with 2% FBS (Gibco, cat# 10270-106) and ACK solution (Gibco, cat. #A1049201), cells were incubated at RT for 5 min. HBSS (13 ml) was added and cells were spun down for 5 min, 1000 rpm, 4°C. Pellets were resuspended in 2 ml pre-warmed 0.05% Trypsin-EDTA (Gibco) and incubated for 5 minutes at 37°C. Serum-free DMEM (3 ml, pre-warmed) and 1μg/ml DNAseI was added and after mixing well, 8 ml of DMEM with 10% FCS was added to stop trypsinization. Before antibody staining, cells were filtered through a 40 μm mesh.

The following antibodies were used: EpCAM-PE (1:100, eBioscience, 12-5791-82, clone G8.8), CD49f-FITC (1:100, eBioscience, 11-0495-82, clone GoH3), CD45-Bio (1:100, eBioscience, 13-0451-82, clone 30-F11), CD31-Bio (1:100, eBioscience, 13-0311-81, clone 390), Ter119-Bio (1:100, eBioscience, 13-5921-81, clone TER-119) and Streptavidin-APC (1:200, eBioscience, 17-4317-82). During antibody staining, cells were in 200 μl HBSS supplemented with 10% HF and kept in the dark on ice for 20 minutes. Before sorting, cells were stained with DAPI (1:5000) and filtered through a 50 μm mesh. Cell sorting was performed using a BD FACS Aria III. FITC was excited with a 488 nm laser and emission was filtered using a 530/30 nm bandpass filter. PE was measured using a 561 nm laser and 582/15 nm bandpass filter. DAPI was excited with a 407 nm laser and emission was filtered using a 450/50 nm bandpass filter. APC was measured using a 633 nm laser and 660/20 nm bandpass filter. Cells were sorted with a plate voltage of 2500 V using the 4-Way Purity precision mode. Cells were collected in TRIzol LS and stored at -80°C.

*RNAscope*

*For SuppFig1B:* The RNAscope 2.5 HD Duplex Assay was used for RNA *in situ*. Freshly isolated mammary glands were dissected and fixed for 24 hours in 4% PFA. Samples were dehydrated through ascending grades of ethanol, cleared in Histo-Clear II (National Diagnostics cat. #HS-200) and embedded in paraffin. Sections of 5 *μ*m were cut using a microtome, after which the Formalin-Fixed Paraffin-Embedded (FFPE) Sample Preparation and Pretreatment protocol was followed (Document Number 322452). One adjustment of the pretreatment was the 30x dilution of Protease Plus with PBS. We optimized this protocol for mammary gland sections and did the Target Retrieval Treatment for 15 minutes and Protease Plus Treatment for 30 minutes. Second part of the RNAscope experiments was done according to RNAscope. 2.5 HD Duplex Detection Kit (Chromogenic) User Manual (Document Number 322500-USM). Probes that were used are: Channel 1: *Krt14* (Cat No. 422521) and Channel 2: *Wnt4* (Cat No. 401101-C2). For the negative controls, RNAscope 2-plex Negative Control Probe Mix was used, which contains premix probes for DapB in both channels. Bright field images were taken on a Zeiss LSM 510 META microscope using a 20x objective in combination with an Axiocam HRc.

#### Document S2

**Supplemental File 1: Details on conserved region 1 and 2 in mCRS4/hCRS4 (related to Fig4)**

**Supplemental File 2: Details on generation of the mCRS4 knockout mice (related to Fig6)**

**Supplemental File 3: Details on 4C *Wnt4* promoter viewpoint (related to Fig2)**

#### Supplemental File 1: Details on conserved region 1 and 2 in mCRS4/hCRS4 (related to Fig4)

Query: mCRS4 (mm10), Length: 1218

BLASTN results: Human DNA sequence from clone RP1-163O16 on chromosome 1p35.1-36.13, complete sequence

Sequence ID: AL031279.1 Length: 135628

**Region 1 (contains conserved placental element lod=73)**

Range: 120682 to 120720

Score: 58.1 bits (63), Expect:6e-05,

Identities: 36/39(92%), Gaps:0/39(0%), Strand: Plus/Minus

*Plus strand mm10:*

CTCFL (prediction mm10, MA1102.2)     CTAGGTGGCGCT     >>>

Query 398 mouse 5’ GGGC**CTAGGTGGCGCT**GTGCACACTTGTCTCCCCAGCCC 436 3’ >>> *Wnt4*

|  |  |  |  |  |  |  |  |  |  |  |  |  |  |  |  |  |  |  |  |  |  |  |  |  |  |  |  |  |  |  |  |  |  |  |

Subjct 120720 human GGGCCTAGGTGGCGATGTGCACACTCGTCTCCCCAGTCC 120682

lod=73 GGGCCTAGGTGGCGCTGTGCACACT chr4:137,164,639-137,164,663

*Minus strand hg38:*

CTCF (prediction hg38, MA1929.2) **CTGCACTGCC**TTCCTTGGGG**CCTAGGTGGCG** >>>

Human 5’ CTGCACTGCCTTCCTTGGGGCCTGGTGGCGATGT 3’ >>> *WNT4*

|  |  |  |  |  |  |  |  |  |  |  |  |  |  |  |  |

Mouse conservation TGTGCTTCCTGCATCTGGGC**CTAGGTGGCGCT**GT

CTCFL (prediction mm10, MA1102.2) **CTAGGTGGCGCT** >>>

**Region 2**

```
Range: 119988 to 120160

Score:122 bits (134), Expect:2e-24,

Identities:133/173(77%), Gaps:3/173(1%), Strand: Plus/Minus

Query  887     TTTATGTCCTGAGCCATTTCCAGCCCAGGT-GAGGCCTTTCTTGCAGAGCCCAGGGG-CA  944
               ||| |||||| ||| |||| |||| ||  | || ||  ||  | ||| |||| |||| ||
Sbjct  120160  TTTGTGTCCTCAGCAATTTTCAGCACAATTTGAAGCTGTTTCTACAGGGCCCCGGGGGCA  120101

GRHL MA1105.3                      AACAGGTT
Query  945     GAAAGAGAAAAATGTGGTCCAACCAGTTACCCGATGTAGTAATAATAACGATAACAGCAA  1004
                  |||||||||||  ||||||||||||| | ||||||||||||||||  |||||| |||
Sbjct  120100  ACCAGAGAAAAATGCAGTCCAACCAGTTATCTGATGTAGTAATAATAAAAATAACAACAA  120041
Lod=62               GAAAAATGTGGTCCAACCAGTTACCCG
Lod=37                                           AGTAATAATAACGATAACAGCAA

Query  1005    CTCCCCTTT-GGTCTGTGTCCCAGCCTCTCGTCTGCATCACCTCAAGTGTCCT  1056
               ||||| |||  ||||   | ||||||| ||  |||||| ||||||  ||||||
Sbjct  120040  CTCCCATTTCTGTCTAGTTTCCAGCCTTTCACCTGCATTACCTCATTTGTCCT  119988
Lod=37        CTC
```

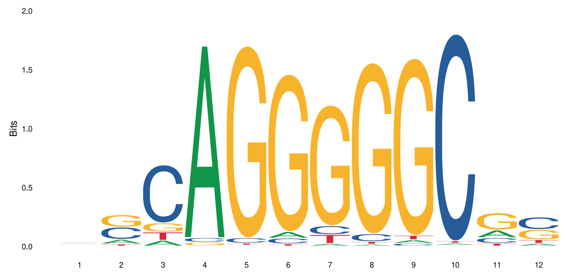

MA1102.2 (CTCFL)

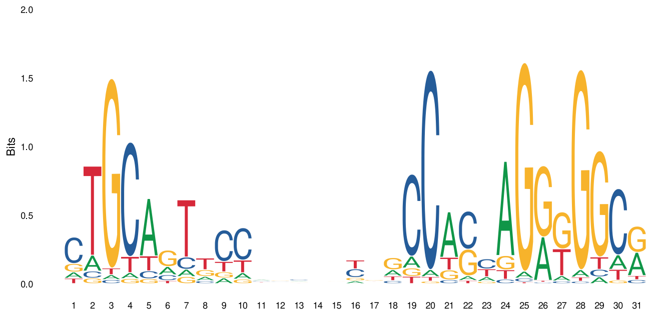

MA1929.2 (CTCF)

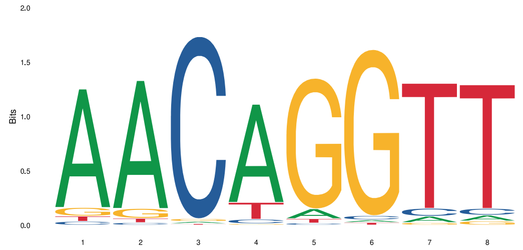

MA1105.3 (GRHL)

#### Supplemental File 2: Details on generation of the mCRS4 knockout mice (related to Fig6)

Overview of the mCRS4 region with annotations (conserved region 1 and 2, Gm13003 TSS, CTCF and GRHL binding sites) and location of the gRNAs and mCRS4 knockout alleles.

Exported from Snapgene.

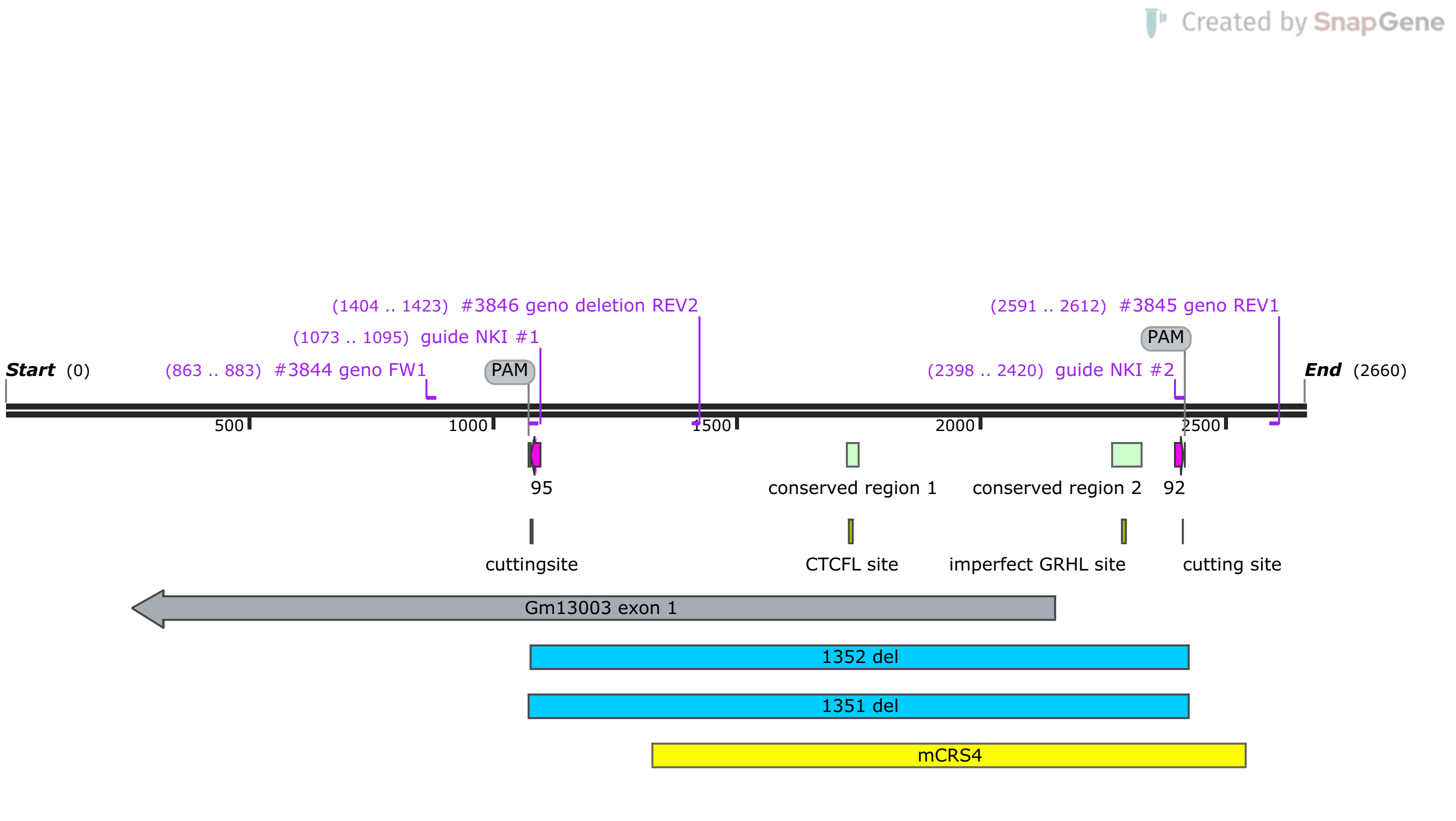

**mm10 coordinates of mCRS4: chr4:137164242-137165457**
2660 bp

#### Supplemental File 3: Details on 4C *Wnt4* promoter viewpoint (related to Fig2)

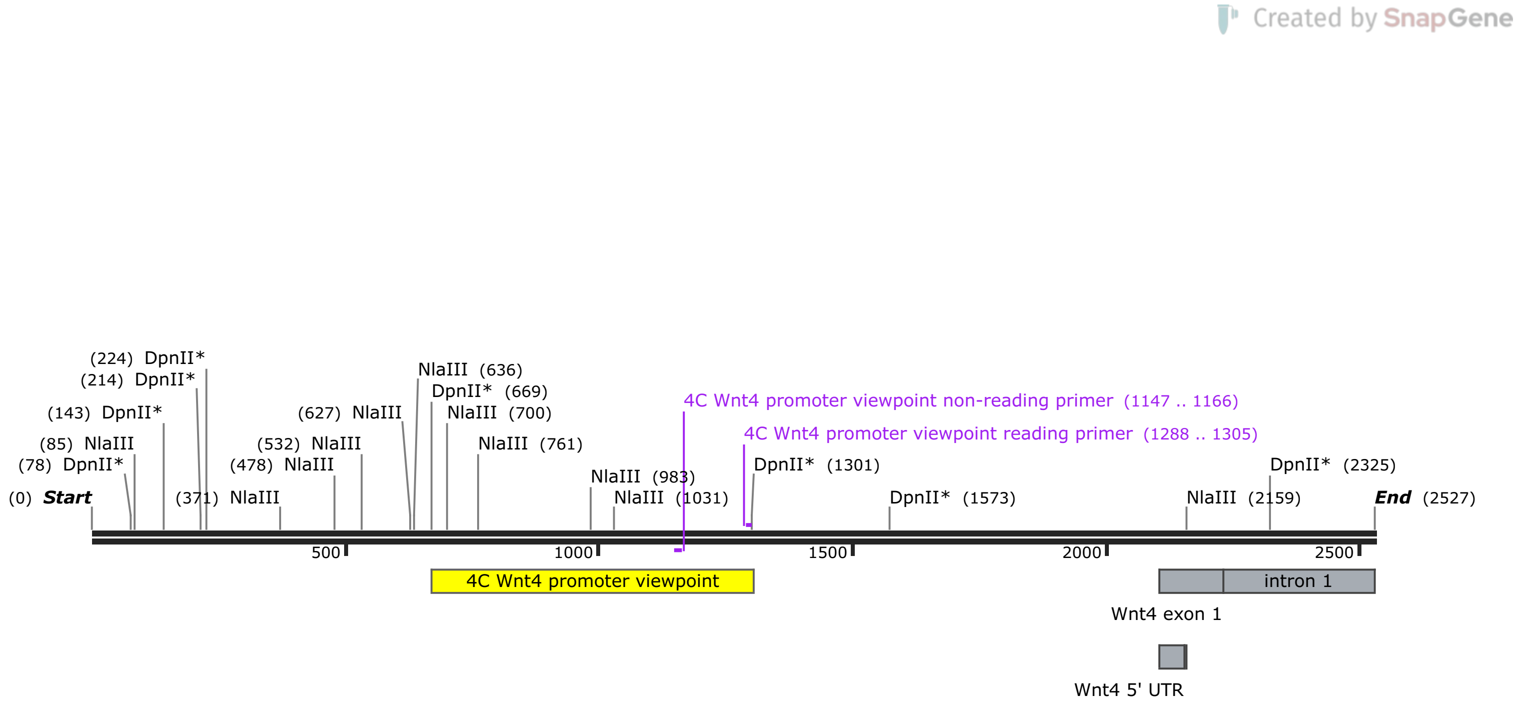

**mm9 DNA for 4C Wnt4 viewpoint design**
2527 bp

#### Document S3

**Supplementary Table 1. Coordinates and conservation of mCRS and hCRS sequences**

**Supplementary Table 2. Mendelian ratios of mCRS4 knockout mice at weaning age**

**Supplementary Table 1. Coordinates and conservation of mCRS and hCRS sequences**
**mCRS/hCRS**: Enhancers were identified as indicated in the main text and numbered from left to right based on their genomic location. Coincidentally, conserved mCRS4 and hCRS4 are fourth in the murine (mCRS1-12) and human (hCRS1-6) sequence. The location of Wnt4/WNT4 is indicated (grey rows). **Coordinates**: Summary of genomic coordinates of mCRS1-12 (top, mm9 and mm10) and hCRS1-6 (bottom, hg19 and hg38). **Overlap with known ENCODE elements**: Indicated where relevant for mouse (mm10) and human (hg38). PLS = promoter like signature, pELS = proximal enhancer like signature, dELS = distal enhancer like signature. n.a. = not annotated. **Sequence conservation**: Results of a BLAST search with the mCRS (or, vice versa the hCRS) sequence on the left and the corresponding hit in the human genome (hg19) (or, vice versa, in the mouse genome, mm10). While conservation exists at the DNA sequence level for multiple mCRS enhancers, only mCRS3, mCRS4 and mCRS6 are also picked up independently via de novo enhancer discovery as conserved human hCRS enhancers (hCRS6, hCRS4 and hCRS1, respectively).

| mCRS | mCRS coordinates (mm9) | mCRS coordinates (mm10) | Overlapping ENCODE elements (mm10) | mCRS sequence conservation (BLAST to hg19) |
|---|---|---|---|---|
| 1 | chr4:136632929-136633690 | chr4:137077015-137077776 | EM10E0773631 (dELS) | chr1:22740837-22741100 |
| 2 | chr4:136653244-136654253 | chr4:137097326-137098339 | EM10E0773635 (dELS) | chr1:22714405-22714515<br>chr1:22714586-22714667 |
| 3 | chr4:136677356-136678081 | chr4:137121443-137122168 | EM10E0773644 (dELS) | chr1:22674972-22675226 |
| 4 | chr4:136720156-136721374 | chr4:137164242-137165457 | EM10E0773650 (PLS)<br>EM10E0773649 (pELS) | chr1:22601380-22602611 |
| 5 | chr4:136765435-136766646 | chr4:137209519-137210735 | n.a. | **not detected** |
| 6 | chr4:136787142-136788282 | chr4:137231229-137232371 | EM10E0773664 (dELS)<br>EM10E0773666 (dELS) | chr1:22531487-22531697 |
| 7 | chr4:136811380-136812040 | chr4:137255464-137256127 | EM10E0773670 (dELS) | **not detected** |
| 8 | chr4:136819669-136820374 | chr4:137263755-137264459 | n.a. | chr1:22486509-22486744 |
| *Wnt4* | chr4:136833550-136852694 | chr4:137277489-137299726 | | |
| 9 | chr4:136835141-136835879 | chr4:137279224-137279963 | n.a. | chr1:22467165-22467657 |
| 10 | chr4:136837266-136838426 | chr4:137281350-137282513 | n.a. | chr1:22465705-22465879 |
| 11 | chr4:136842973-136843761 | chr4:137287060-137287845 | n.a. | chr1:22459188-22459398 |
| 12 | chr4:136863009-136863969 | chr4:137307093-137308056 | EM10E0773691 (dELS) | chr1:22433396-22433929 |

| hCRS | hCRS coordinates (hg19) | hCRS coordinates (hg38) | Overlapping ENCODE elements (hg38) | hCRS sequence conservation (BLAST to mm10) |
|---|---|---|---|---|
| *WNT4* | chr1:22443806-22469590 | | | |
| 1 (mCRS6-like) | chr1:22531184-22532181 | chr1:22204691-22205688 | n.a. | chr4:137231684-137231893 |
| 2 | chr1:22583411-22585067 | chr1:22256918-22258574 | EH38E1326868 (dELS)<br>EH38E1326869 (dELS) | chr4:137182368-137183425 |
| 3 | chr1:22585640-22586636 | chr1:22259147-22260143 | EH38E1326871 (dELS)<br>EH38E1326872 (dELS) | **not detected** |
| 4 (mCRS4-like) | chr1:22601504-22603057 | chr1:22275011-22276564 | EH38E1326895 (dELS)<br>EH38E1326896 (dELS)<br>EH38E1326897 (dELS) | chr4:137164639-137164677<br>chr4:137165128-137165297 |
| 5 | chr1:22622582-22624241 | chr1:22296089-22297748 | EH38E1326913 (dELS)<br>EH38E1326914 (dELS)<br>EH38E1326915 (dELS) | **not detected** |
| 6 (mCRS3-like) | chr1:22674614-22675610 | chr1:22348121-22349117 | EH38E1326941 (dELS) | chr4:137121785-137121955 |

**Supplementary Table 2. Mendelian ratios of mCRS4 knockout mice at weaning age**
Heterozygous crosses were performed for two independent mCRS4 knock-out lines (mCRS4Δ1 and mCRS4Δ2). At weaning, no statistically significant differences were observed in either the male:female ratio ($P_{sex}$, Chi-square test) or the distribution of wildtype, heterozygous or homozygous animals ($P_{geno}$, Chi-square test).
This table complements Fig6.

| Parental cross | Offspring at weaning | | | | | | | | | |
|---|---|---|---|---|---|---|---|---|---|---|
| | No. pups | Sex | No. Expected | No. Observed | $P_{sex}$ | Genotype | | No. Expected | No. Observed | $P_{geno}$ |
| Wnt4 $^{+/mCRS4\Delta1}$ x Wnt4 $^{+/mCRS4\Delta1}$ | 142 | Male | 71 | 66 | 0.395 (ns) | WT | Wnt4 $^{+/+}$ | 16.5 | 14 | 0.562 (ns) |
| | | | | | | HET | Wnt4 $^{+/mCRS4\Delta1}$ | 33 | 32 | |
| | | | | | | HOM | Wnt4 $^{mCRS4\Delta1/mCRS4\Delta1}$ | 16.5 | 20 | |
| | | Female | 71 | 76 | | WT | Wnt4 $^{+/+}$ | 19 | 17 | 0.779 (ns) |
| | | | | | | HET | Wnt4 $^{+/mCRS4\Delta1}$ | 38 | 41 | |
| | | | | | | HOM | Wnt4 $^{mCRS4\Delta1/mCRS4\Delta1}$ | 19 | 18 | |
| Wnt4 $^{+/mCRS4\Delta2}$ x Wnt4 $^{+/mCRS4\Delta2}$ | 145 | Male | 73 | 77 | 0.464 (ns) | WT | Wnt4 $^{+/+}$ | 19.25 | 14 | 0.145 (ns) |
| | | | | | | HET | Wnt4 $^{+/mCRS4\Delta2}$ | 38.5 | 47 | |
| | | | | | | HOM | Wnt4 $^{mCRS4\Delta2/mCRS4\Delta2}$ | 19.25 | 16 | |
| | | Female | 73 | 68 | | WT | Wnt4 $^{+/+}$ | 17.25 | 10 | 0.138 (ns) |
| | | | | | | HET | Wnt4 $^{+/mCRS4\Delta2}$ | 34.5 | 40 | |
| | | | | | | HOM | Wnt4 $^{mCRS4\Delta2/mCRS4\Delta2}$ | 17.25 | 18 | |
